# Mesoscale molecular architecture of the human striatum across cell types and lifespan

**DOI:** 10.64898/2026.03.04.709715

**Authors:** Andrew W. Kraft, Matthew Lee, Nirmala Rayan, Haoyuan Gao, Julianna Milidantri, Charles Vanderburg, Karol Balderrama, Naeem Nadaf, Vipin Kumar, Katelyn Flowers, Emily Finn, Matthew Shabet, Ezra Muratoglu, Olivia Yoo, Khalid Shakir, James Nemesh, Steven Burger, Sadie Drouin, Olivia Catalini, Mukund Raj, Abir Mohsin, Nikita Budnik, Lucas Reese, Steven A. McCarroll, Kiku Ichihara, Evan Z. Macosko

## Abstract

The human striatum is a central hub for diverse motor, cognitive, and affective behaviors, yet it lacks obvious cytoarchitectural boundaries defining functional territories. Here, we uncover a robust and molecularly defined mesoscale architecture in the human striatum. Using Slide-tags, a scalable single-nucleus spatial transcriptomics technology, we profiled 1.1 million cells across the striatum of 19 postmortem donors. Our data uncover a natural subdivision into six zones, each defined by distinct medium spiny neuron (MSN) populations and coordinated neuron-astrocyte signaling. Dorsal zone MSNs exhibit higher expression of synaptic remodeling and plasticity genes, while ventral zones are enriched for semaphorin, chaperone, and hedgehog signaling. Imputing these identities onto a larger 131-donor RNA-seq cohort, we find dorsal zones show greater age-related transcriptional changes, and spatial zonation attenuates with age. This atlas provides a mesoscale molecular definition of striatal anatomy, linking cell identity to functional specialization and aging susceptibility.

## Introduction

The human striatum is the basal ganglia’s primary input nucleus, essential for integrating cortical and thalamic information to drive a wide range of motor, cognitive, and affective behaviors^1^. Anatomical tract-tracing studies in non-human primates have established that this structure is organized into topographical boundaries: a “limbic” domain in the ventral striatum, an “associative” domain in the caudate region, and a “sensorimotor” domain in the putamen^2,3^. Similarly, viral tracing studies in rodents have defined mesoscale domains based upon patterns of corticostriatal input^4,5^. This functional topography further distinguishes regions of selective vulnerability–from the motor deficits of Parkinson’s disease in the putamen^6^, to the affective dysregulation of addiction in the nucleus accumbens^7^. However, unlike the cerebral cortex, where functional boundaries are often demarcated by clear cytoarchitectural transitions, these striatal territories lack obvious anatomical borders in human tissue^8^. Immunohistochemical studies have demonstrated regionally restricted expression of a limited number of genes across the striatum. However, we lack a comprehensive, spatially resolved molecular framework capable of robustly and consistently defining these domains, leaving a critical gap between our understanding of striatal circuits and the molecular identity of the cells that compose them.

Deepening this complexity is the organization of the striatum at smaller scales. The medium spiny neurons (MSNs, also described in many studies as striatal projection neurons or SPNs)--the principal neurons of the striatum–are organized into two compartments: the matrix, constituting the majority of cells (∼80%), and the “patches,” or “striosomes”--collections of MSNs, 300-600 µm in diameter, that possess distinct molecular and circuit identities^9,10^. Matrix and striosomal MSNs are both further subdivided by their “pathway” designation: D1 dopamine receptor-expressing MSNs primarily project directly to the substantia nigra, while D2 dopamine receptor-expressing MSNs primarily project to the globus pallidus. Yet, the governing principles that relate these microscale partitions to one another–and to the broader functional territories of the striatum–remain incompletely understood.

Molecular profiling by single-cell and spatial genomics offers a potential opportunity to unify these molecular, cellular, and functional domains of study. In mice, single-cell and single-nucleus RNA sequencing (snRNA-seq) has discovered new continuous and discrete axes of variation in MSNs beyond the longstanding distinctions based on connectivity (direct vs. indirect) and compartment (matrix vs. patch/striosome)^11–13^. Spatially resolved studies in the rodent and nonhuman primate models have begun to assemble maps defining patch, matrix, and “exopatch” domains alongside larger territories^14,15^. A key remaining question is whether these distinct scales of organization—molecular phenotypes, patch-matrix compartments, and larger territorial domains—are independent features, or whether they adhere to a unifying, hierarchically organized framework.

Fully resolving the molecular topography of the human striatum requires a technology that can simultaneously perform genome-wide measurements of gene expression to identify subtle variation in cell types, while covering centimeter-sized spatial scales across multiple postmortem samples. This is critical to identifying conserved, consistent organizational principles that traverse the full neuroanatomical extent of the structure. We recently developed Slide-tags, a technology that analyzes large tissue areas at single-nucleus resolution^16,17^. Slide-tags transfers spatial barcodes from a large-area array directly into the nuclei of intact tissue sections prior to dissociation. This allows the recovery of high-quality single-nucleus transcriptomes while also ascertaining a cell’s precise spatial coordinates.

Here, we apply this technology to transcriptionally profile the human striatum from 19 donors, across 1.1 million cells. By performing our spatial assay on many donors, we identified reproducible cell states and cellular compositions across the striatum that were consistent across all human donors in the analysis. Our analyses uncovered a zonation pattern that was conserved across all 19 donors, present most prominently in the matrix MSNs but also found in all other MSN subtypes and in astrocytes. We detail the cellular composition of each zone, how each cell type specializes across zones, and examine how zonal identity changes with normal aging.

## Results

### Slide-tags defines the compartmental cytoarchitecture of the human striatum

To systematically uncover the cellular cytoarchitecture of the human striatum, we scaled Slide-tags to enable the spatial profiling of tissue specimens spanning several square centimeters (**Methods)**. Deploying these arrays on pre-commissural striatal sections containing the head of the caudate, putamen, and nucleus accumbens, from 19 postmortem donors with broad demographic representation (**Figure 1A**, **S1A, Table S1**), we recovered between 46,066 and 151,068 snRNA-seq profiles across 7 cm^2^ of tissue per donor (**Table S1**). We successfully spatially localized a total of 1,102,999 nuclei, maintaining a negligible false-placement rate as validated by anatomical white-matter fiducials (**Figure 1B, Figure S1B,C).** Transcript capture sensitivity matched that of standard snRNA-seq performed on a larger reference cohort (**Figure S1D)**. At the class level, we recovered and mapped all canonical striatal populations, including MSNs, interneurons, astrocytes, oligodendrocytes, vascular cells, and microglia (**Figure S1F**). Label transfer from a recent basal ganglia reference taxonomy^18^ confirmed the spatial placement of 10 molecularly defined MSN populations, 11 interneuron types, and 13 non-neuronal types (**Figure S1E,F**).

**Figure 1.**
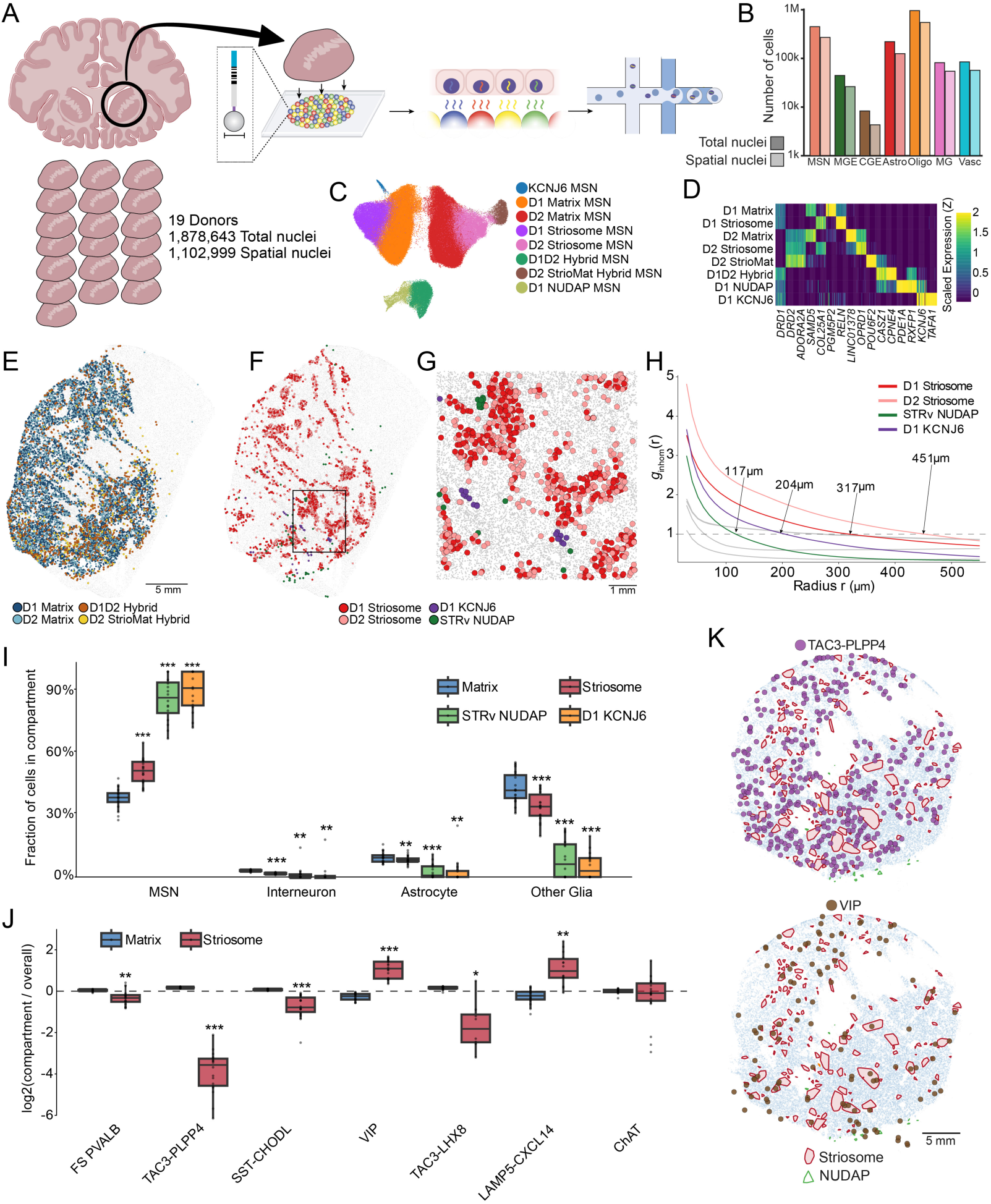
Slide-tags profiling reveals the compartmental micro-scale structure of the human striatum. **A)** Experimental workflow for Slide-tags spatial profiling of 19 postmortem human striatal sections. **B)** Total nuclei recovered per major cell class. **C)** UMAP embedding of medium spiny neuron (MSN) single-nucleus expression profiles, colored by subtype. **D)** Normalized marker gene expression defining MSN subtypes. **E–G)** Spatial distributions of diffusely distributed **(E)** and highly compartmentalized **(F)** MSN subtypes across the tissue, with a magnified view of the clustered populations **(G)**. **H)** Inhomogeneous pair correlation (g_inhom(r)) estimating cluster radii for MSN subtypes. Colored lines denote locally compartmentalized types; gray lines denote diffuse types. Cluster radii are defined by the first (g_inhom(r) = 1) intercept, where local neighbor density drops to the background expected value. **I)** Major cell class composition within spatially defined compartments. **J)** Log2 fold enrichment of interneuron subtypes in Striosomes versus Matrix. Asterisks denote Benjamini-Hochberg corrected paired Wilcoxon signed-rank tests comparing each non-Matrix compartment to the Matrix across donors (*p < 0.05, **p < 0.01, ***p < 0.001). **K)** Spatial distributions of TAC3-PLPP4 (top) and VIP (bottom) interneurons in a representative donor. Convex hulls delineate compartments defined by DBSCAN clustering.

We next focused on micro-scale spatial architecture of the MSN subsets. The two principal compartments of the striatum displayed their canonical topographies: Matrix neurons were distributed evenly throughout the structure, while Striosome MSNs formed discrete, spatially restricted patches visible in every donor (**Figure 1C-E, S1G**). Both compartments were further subdivided by pathway identity into *DRD1*-expressing (D1 Direct) and *DRD2*-expressing (D2 Indirect) populations (**Figure 1D**). Beyond these cardinal populations, we mapped rarer, more molecularly distinct MSN subtypes present in the reference taxonomy, including the Hybrid (or “Eccentric”), STRv NUDAP and StrioMat MSNs (**Figure 1D,E,F,G**). Furthermore, we identified a sparse, molecularly distinct MSN population–marked by the selective expression of *KCNJ6* and *TAFA1*– that was not present in the reference (**Figure 1D,F,G**). While the Matrix, StrioMat, and Hybrid populations were diffusely intermingled throughout the tissue, the STRv NUDAP and D1 KCNJ6 subtypes exhibited clustered distributions localized primarily in the nucleus accumbens and the ventrolateral base of the putamen, suggesting they form distinct spatial patches akin to striosomes (**Figure 1F,G**).

To systematically quantify this organization, we applied an inhomogeneous pair correlation function (**Methods)**. By expanding the radial search space from each cell and normalizing against an MSN-wide or type-specific null model (**Methods**), we determined which MSN types displayed significantly clustered local architecture, and estimated the maximal physical extent of each of these structural compartments. This analysis confirmed that the D1 and D2 Striosome MSNs, along with the STRv NUDAP and D1 *KCNJ6* populations, exhibit significant spatial compartmentalization, whereas the Matrix and Hybrid MSNs do not (**Figures 1G, S1H**). The metric estimated the average radii of the striosomal compartments to be 317 to 451 µm, consistent with neurochemical estimates^10,19^, while the D1 *KCNJ6* and STRv NUDAP compartments formed considerably tighter domains (204 µm and 117 µm, respectively, **Figure 1H**).

To understand how these micro-compartments shape local cellular neighborhoods, we spatially clustered the tissue from each donor to segment the gray matter into Matrix, Striosome, NUDAP, and *KCNJ6* territories (**Methods**). Consistent with previous observations across species, interneurons as a class were depleted from striosomes, and were nearly absent from the STRv NUDAP and D1 *KCNJ6* clusters (**Figure 1I**). Intriguingly, while the depletion of PV and SST interneurons from striosomes has been noted^6,20^, our human data revealed that the TAC3-PLPP4 interneuron subtype is the most highly Matrix-specific (−4.08 +/- 0.24 log2 fold change). By contrast, rarer interneuron populations—most notably the VIP and LAMP5 subtypes—were preferentially enriched in the striosomes (**Figure 1J,K**). Collectively, our data establish a comprehensive molecular and physical definition of the striatum’s micro-scale architecture, anchored by four highly compartmentalized, spatially distinct cellular domains.

### Consistent mesoscale territories emerge from unsupervised clustering of MSNs and astrocytes

To explore larger scale spatial organization within MSN types, we first performed unsupervised clustering specifically on the D1 Matrix MSNs (**Methods**). Dimensionality reduction revealed that these cells form a continuous, ring-like topology in gene expression space (**Figure 2A**).

**Figure 2.**
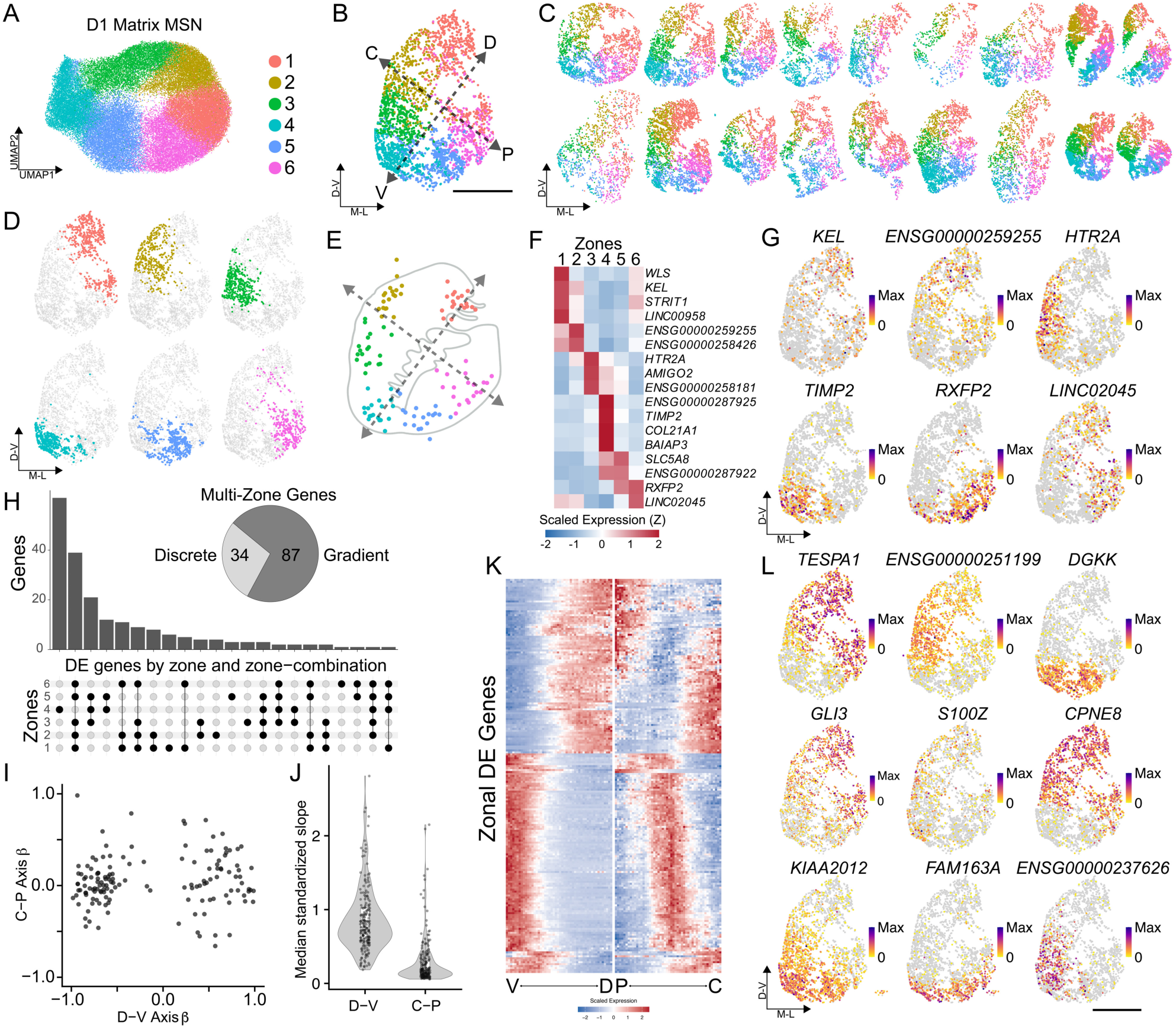
Molecular mesoscale divisions of D1 Matrix neurons across the human striatum. **A)** UMAP embedding of all D1 Matrix MSNs identified from the 19 profiled donors, colored by unsupervised cluster identity. **B-D)** Spatial distribution of D1 Matrix MSNs shown for **(B)** a representative donor, **(C)** all 19 donors, and **(D)** a representative donor with each of the six zones plotted individually. Anatomical orientations: C, caudate; P, putamen; D, dorsal; V, ventral. **E)** Centroid positions of each zone across all donors, aligned in a transformed common coordinate framework. This atlas space is defined by a dorsal-ventral (D-V) axis—anchored by the centroids of Zones 1 and 4—and an orthogonal caudate-putamen (C-P) axis. **F-G)** Gene expression heatmap **(F)** and corresponding spatial plots **(G)** of discrete marker genes defining the six zones. **H**) UpSet plot (left) detailing differentially expressed genes (DEGs) shared across individual zones or combinations of zones. The pie chart (right) classifies multi-zone DEGs as either "discrete" (equivalent expression across positively expressing zones) or "gradient" (varying expression between positively expressing zones). **I)** Expression gradients for all DEGs identified in **(H)**, plotting the slope of expression along the projected D-V axis (x-axis) versus the C-P axis (y-axis). **J)** Quantification of expression slopes, representing the median value across donors for each DEG. **K)** Heatmap illustrating the expression of zonal DEGs smoothed across the D-V and C-P spatial axes. **L**) Spatial distribution of representative gradient and discrete multi-zone DEGs, visualized in a single donor.

When the positions of the emergent clusters were mapped back onto the physical tissue coordinates across the 19 donors, we observed that this abstract “ring” in expression space corresponded to a remarkably consistent, zonated distribution in physical space (**Figure 2B-D**). Upon transformation into a shared coordinate framework (**Methods**), the cluster centroids aligned tightly across all 19 donors, indicating a highly conserved spatial architecture (**Figure 2E**). Crucially, these six molecular “zones” did not fully obey classical anatomical subdivisions. Most notably, Zone 1 bridged the dorsolateral caudate and the putamen, suggesting a shared molecular identity across these anatomically distinct structures, while Zones 4 and 5 were confined to the ventral base of the caudoputamen and nucleus accumbens.

Given that cluster resolution is a parameter-dependent choice, we sought a principled approach to define the biologically optimal granularity of these zones. We prioritized two criteria: spatial coherence (quantified via kernel density estimation) and cross-donor reproducibility. Titration of the resolution parameter revealed that k=6 was the maximum granularity at which clusters remained spatially contiguous and non-overlapping while maintaining robust representation across all donors (**Figure S2A-D**). This establishes that a six-zone parcellation represents the finest-grained molecular definition of D1 matrix cells supported by the data.

To investigate the molecular basis of these zones, we performed differential gene expression analysis, identifying a total of 199 markers distributed in unique combinations across the zones. Of these, 78 were zone-specific, with every zone marked by at least one discrete gene (**Figures 2F-H**). The remaining 121 genes were differentially expressed across varied combinations of multi-zone blocks, further distinguishing zonal identities (**Figure 2H**). In particular, Zone 4, located at the tip of the ventral striatum, was most distinct from the other zones, and in general, zones next to each other spatially showed the strongest gene expression commonalities (**Figure S2E**). We therefore examined whether many gene expression differences existed along a continuum in space. To evaluate this, we constructed two splines, one running along the dorsal-ventral axis, and another running perpendicular along the medial-lateral axis (**Methods**). We then tested the correlation of each differentially expressed gene along this axis. Most genes showed a dependency on the dorsal-ventral axis, while only a handful of genes varied along the medial-lateral axis (**Figures 2I-L**). Striatal zones are thus primarily defined by their dorsal-ventral position, with a smaller set of genes discriminating them from each other in a discrete fashion.

We then examined whether this six-zone molecular architecture is conserved along the anterior-posterior (A-P) axis. We applied Slide-tags to the posterior striatum of two human donors (**Figure S2F, Methods**). Integrative analysis of D1 Matrix MSNs revealed that the posterior striatum is compositionally enriched for Zones 1, 2, and 6 (**Figure S2G-I**), and most crucially, contains no novel molecular clusters not observed in the more anterior sections. Zonal marker expression remained highly concordant regardless of A-P position (**Figure S2J**), indicating that this six-zone framework defines the full longitudinal extent of the sampled striatum. Additionally, to determine if this cytoarchitecture extends beyond humans, we profiled three serial coronal striatal sections from an adult macaque (*Macaca fascicularis*, **Figure S2K**). Unsupervised clustering of the 7745 recovered macaque D1 Matrix neurons revealed a spatial topology remarkably homologous to the human tissue (**Figure S2L**), with zonal centroids aligning well with those of the human (**Figure S2M**). Our molecular parcellation thus represents a consistent and evolutionarily conserved structural feature of the primate basal ganglia.

We next investigated whether this zonal architecture extends across all MSN populations. Unsupervised clustering of D2 Matrix MSNs revealed a remarkably similar ring-like topology in UMAP space (**Figure 3A**) and near-identical spatial localizations (**Figures 3B**), with *k*=6 confirmed as the optimal granularity (**Figures S2A-C**). Spatial boundaries and transcriptional signatures were highly conserved across pathways: corresponding D1 and D2 zones exhibited >75% spatial overlap (**Figures 3C, S3B**), while D2 marker genes displayed similar combinatorial logic, dorsal-ventral expression gradients (**Figure S3D**), and strong correlations with their D1 counterparts (**Figures S3E,F**). Applying this unsupervised approach to the rarer striosome and Hybrid (eSPN) populations (**Methods**) yielded a similar continuous topology; however, likely owing to their comparative sparsity, these cells reliably resolved into only four cross-donor coherent territories (**Figures 3D-J**). Nevertheless, these four domains adhered to the broader matrix contours—Matrix Zone 1 aligned with the non-matrix Zone D, Zones 2 and 3 merged into Zone C, Zone 4 mapped to Zone V, and Zones 5 and 6 unified into Zone P (**Figures 3F,H; S3H**)—indicating that all MSN subtypes follow a consistent spatial cytoarchitecture.

**Figure 3.**
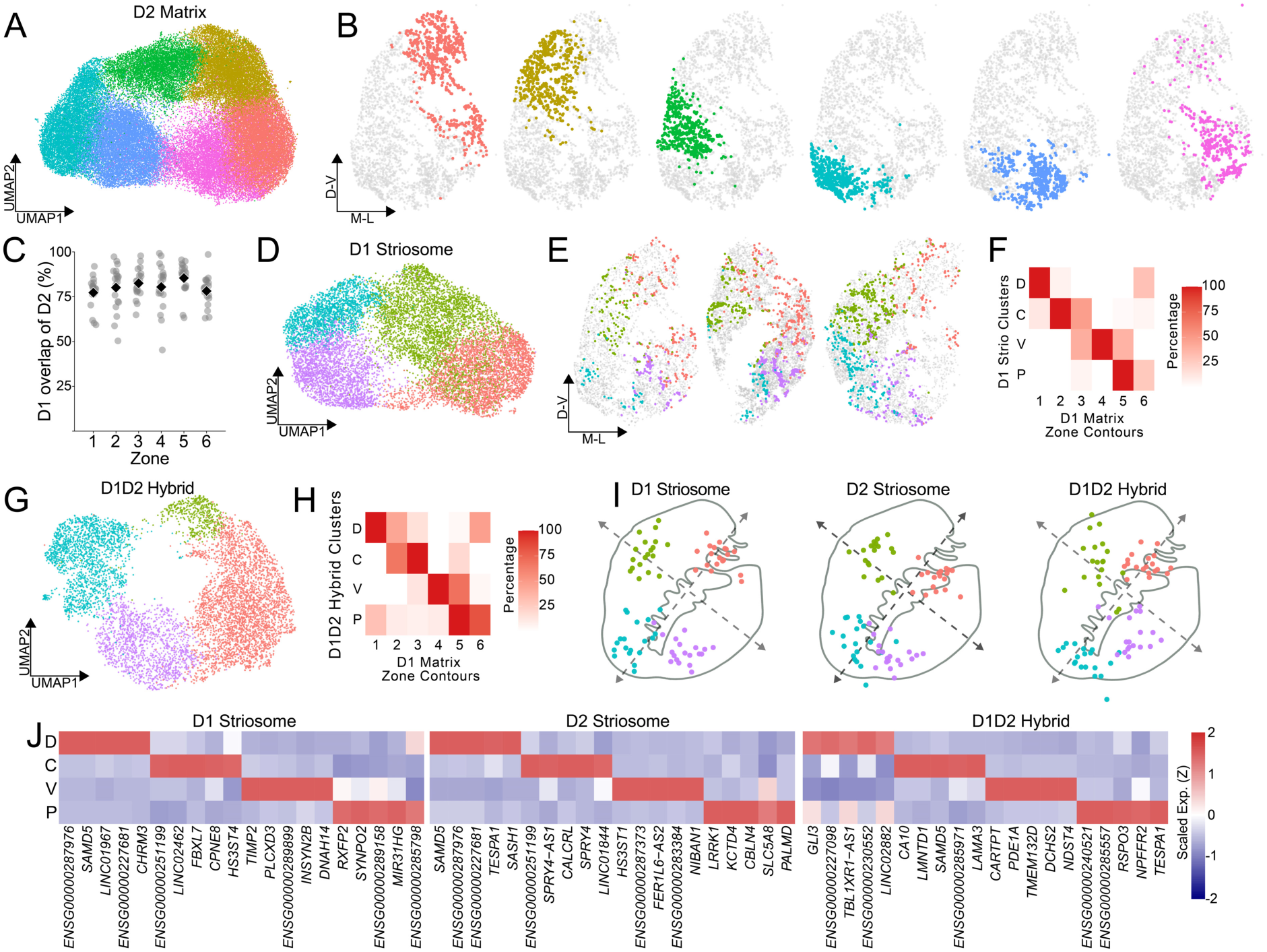
Molecular zones are conserved across diverse MSN subtypes. **A-B)** Unsupervised clustering of D2 Matrix neurons from all 19 donors, visualized in (**A**) UMAP space and (**B**) physical coordinates of a representative donor. **C)** Spatial overlap of kernel density estimation (KDE)-defined contours (see **Methods**) between the six zones identified independently in D1 and D2 matrix neurons. **D-E**) Unsupervised clustering of D1 Striosome neurons, visualized in **(D)** UMAP space and **(E)** physical spatial coordinates across three representative donors. **F**) Spatial contour overlap between D1 Striosome clusters (rows) and D1 Matrix zones (columns). **G-H**) UMAP embedding **(G)** and spatial contour overlap with D1 Matrix zones **(H)** for D1/D2 Hybrid MSNs. **I**) Spatial alignment of cluster centroids across all 19 donors for D1 Striosome (left), D2 Striosome (middle), and D1/D2 Hybrid (right) populations, demonstrating cross-donor spatial conservation. **J)** Heatmap of normalized expression for zone-specific marker genes across the evaluated MSN populations.

We next asked whether this spatial architecture is exclusive to neurons, or if it also organizes striatal glia. We first performed unsupervised clustering across six major non-neuronal classes. With the sole exception of endothelial cells, every non-neuronal population exhibited a fundamental transcriptional dichotomy distinguishing the gray matter from the surrounding white matter tracts (**Figures S4A,B**). Focusing specifically on gray matter astrocytes, subclustering revealed four robust, cross-donor reproducible spatial populations (**Figure 4A-C, Figure S4C,D**). One astrocytic population aligned precisely with MSN Zone 1, a second bridged the associative Zones 2 and 3, a third encompassed the ventral Zones 4 and 5, and the fourth localized to Zone 6 (**Figure S4E**). Defined by the expression of highly discrete spatial markers (**Figure 4D,E**), and conserved across all 19 donors (**Figure S4F**), these findings demonstrate that human striatal zones are not exclusively a neuronal phenomenon.

**Figure 4.**
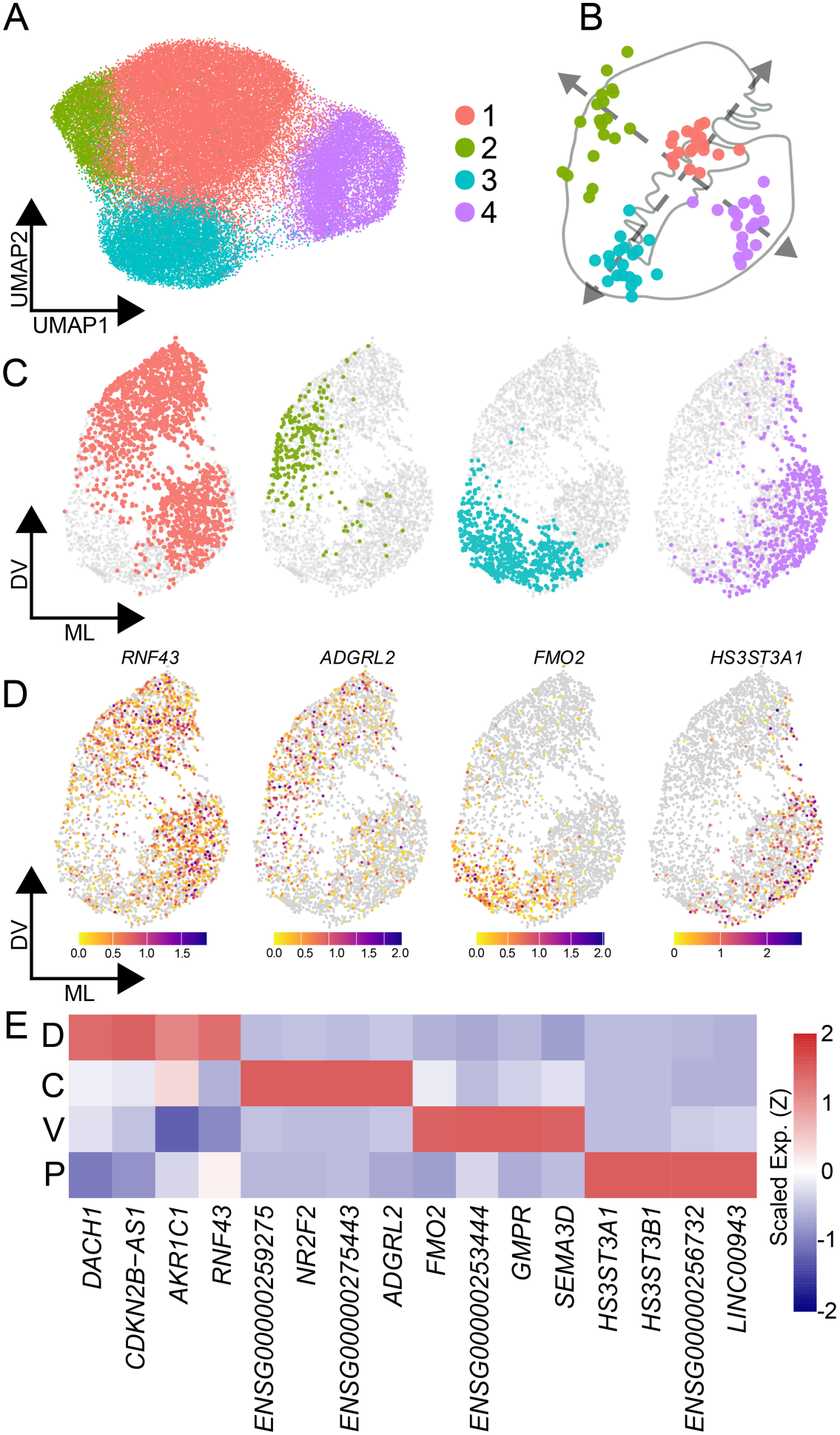
Molecular zones of the human striatum extend to astrocytes. **A–C)** Unsupervised clustering of astrocytes from all 19 donors, visualized **(A)** in UMAP space, **(B)** as aligned spatial cluster centroids across all donors, and **(C)** in the physical spatial coordinates of a representative donor. **D)** Spatial expression plots of representative zone-specific marker genes in astrocytes. **E)** Heatmap of normalized expression for zone-specific marker genes across astrocytic populations.

### Distinct cellular assemblies and pathway biases in striatal zones

Having established the consistency and robustness of striatal zones, we next investigated whether these spatial domains correspond to distinct cellular microenvironments. We assigned each cell, in each donor, to a zonal identity based upon identities of the nearest neighboring D1 matrix MSNs (**Figure 5A**). As expected, the zones in the most dorsal aspect of the striatum, where penetrating white matter fibers are densest (Zones 1, 2, and 6) showed the highest proportion of oligodendrocytes. In contrast, the ventral Zone 4 had the lowest oligodendrocyte proportion and the highest fraction of neurons (**Figures 5B, S5A**), reflecting the distinct biophysical environment of the nucleus accumbens^21^.

**Figure 5.**
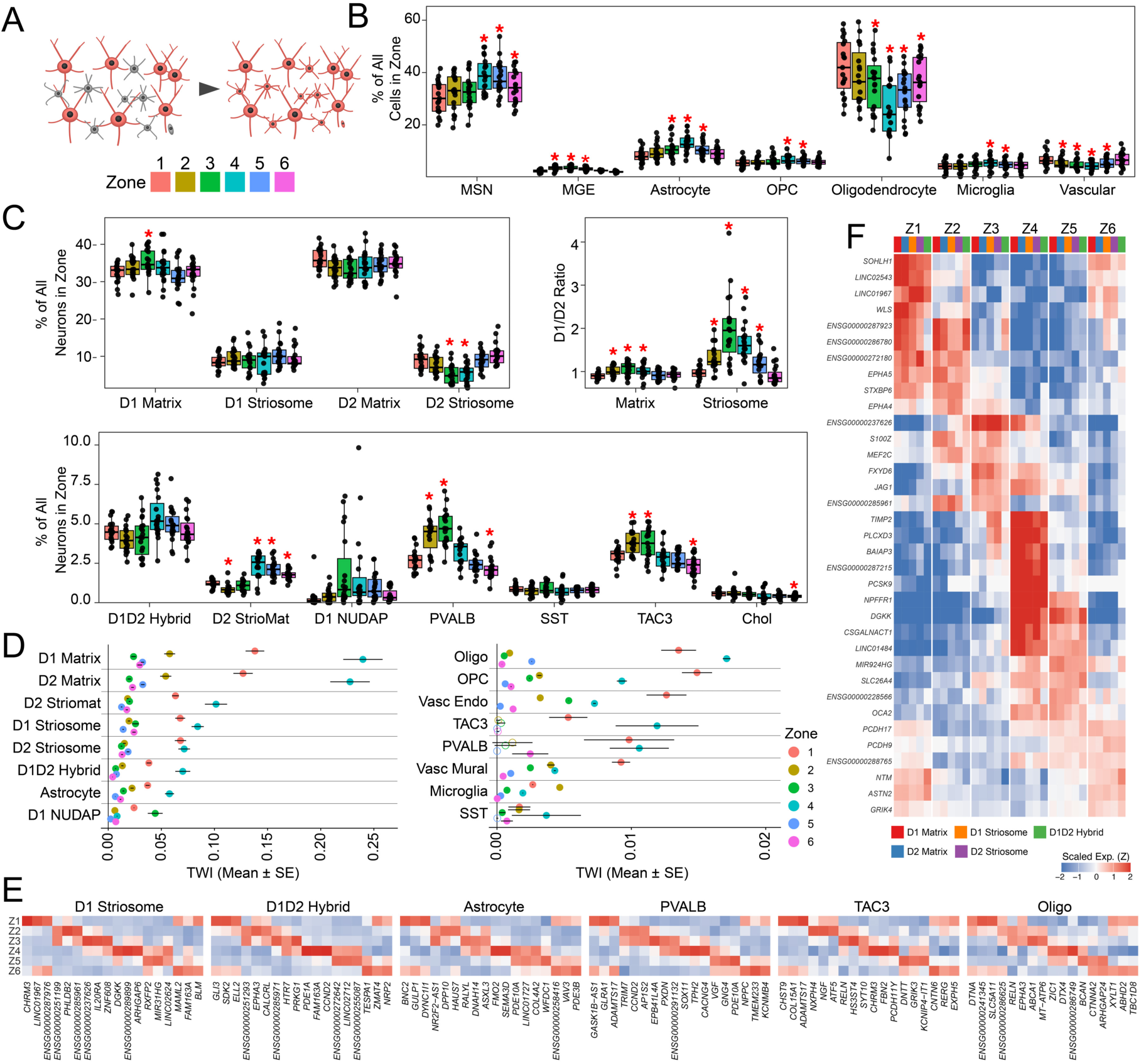
Striatal zones possess distinct cellular architectures and transcriptional specializations. **A)** Schematic illustrating the assignment of zonal identities to all cell types based on their spatial proximity to reference D1 Matrix MSNs. **B)** Proportional representation of major cell classes within each of the six spatial zones. **C)** Fractional composition of matrix and striosome MSN subtypes (upper left) and other neuronal populations (bottom) across zones, alongside the shifting ratio of D1 to D2 MSNs (upper right). **D)** Quantification of Transcriptome-Wide Impact (TWI)—a metric of molecular distinctiveness—for each cell type across the six zones. **E)** Heatmap of normalized expression for representative zone-specific marker genes across D1 Striosome MSNs, D1D2 Hybrid MSNs, Astrocytes, PVALB and TAC3 Interneurons, and oligodendrocytes. **F)** Heatmap of normalized expression for core MSN zone-specific genes that are common across Matrix, Striosome and D1D2 Hybrid MSNs. *p < 0.05, **p < 0.01, ***p < 0.001.

While the canonical model of the basal ganglia posits a balanced opposition between the D1 and D2 pathways, our spatial analysis reveals that this stoichiometry fluctuates across zones. In the caudate-specific Zones 2 and 3, we observed a proportional expansion of D1 Matrix MSNs (**Figure 5C**), resulting in a significant D1 bias within these territories (**Figure 5C**). These findings mirror gradients reported in the rodent striatum, where ‘D2-poor’ zones have been identified in caudal^22^ and ventromedial^23^ subregions. Intriguingly, this pathway bias was not confined to the matrix; the D1:D2 ratio of striosomes strongly covaried with that of the surrounding matrix (**Figure 5C**). This demonstrates that despite their distinct connectivities, both compartments are subject to a shared, zone-specific spatial architecture. Functionally, these data suggest that the associative striatum (Zones 2 and 3) is structurally biased toward D1 pathway signaling, whereas the sensorimotor zones (Zones 1,5, and 6) have a more balanced or D2-skewed architecture. The D1-biased environment in Zones 2 and 3 was accompanied by a coordinated shift in interneuron architecture, specifically the co-enrichment of TAC3 and PVALB interneurons (**Figure 5D**). The co-enrichment of PVALB fast-spiking interneurons is consistent with a compensatory “gain control” mechanism, as they have been shown to provide stronger, more effective feed-forward inhibition onto D1 MSNs than D2 MSNs in many contexts^24^, connecting their proportional increase in zones with higher D1 MSN representation.

Beyond these shifts in cellular composition, we next investigated the magnitude of *molecular* specialization within these assemblies. Using a transcriptome-wide impact (TWI) score—which measures the aggregate shift in gene expression for a given cell type across zones^25^–we found that MSNs and astrocytes exhibit the highest degree of zonal specialization (**Figure 5D**).

Furthermore, Zones 1 and 4 displayed the strongest molecular distinctions relative to the other zones across all cell types, anchoring them as the most transcriptionally polarized compartments within the striatum.

To understand the nature of this specialization, we evaluated the zone-specific marker genes across all cell types. By anchoring our analysis to the matrix MSN spatial coordinates, we achieved higher spatial resolution than unsupervised clustering alone, revealing that Striosome MSNs, D1/D2 Hybrid MSNs, and astrocytes express distinct spatial markers across all six zones. Strikingly, the zonal marker genes were highly correlated across all MSN subtypes, defining a "core MSN zonal gene signature" that overrides individual subtype identities (**Figures 5E,F**). By contrast, the zonal signatures of astrocytes, interneurons, and other glia—including OPCs, whose spatial specificity is only recently being appreciated—were unique and uncorrelated with the MSN signatures (**Figures 5E, S5B-D**). These results establish that human striatal zones profoundly shape the transcriptional state of all resident cell types, driving a shared spatial program in MSNs while inducing distinct, tailored specializations in the interneurons and surrounding glia.

### Molecular specialization within striatal zones

Having defined the cellular architecture of human striatal zones, we next sought to understand the functional implications of these spatial programs. We focused on the Matrix MSNs and astrocytes in Zones 1 and 4, which our transcriptome-wide impact analyses indicated were the most molecularly distinct. Gene Set Enrichment Analysis (GSEA) of the differentially expressed programs revealed clear differences between the dorsal and ventral poles of the striatum (**Figures 6A-I, S6, Table S2**). Zone 1 MSNs displayed significantly higher expression of genes that promote and regulate synaptic plasticity, including *GRID2IP*, *GRIN2A*, and *PTN* (**Figure 6D)**. By contrast, Zone 4 MSNs were enriched for the expression of heat shock proteins and protein chaperones, including *HSPA5*, *HSPA8*, and *HSP90AB1* (**Figure 6D**), suggesting a heightened requirement for proteostatic maintenance in the ventral striatum.

**Figure 6.**
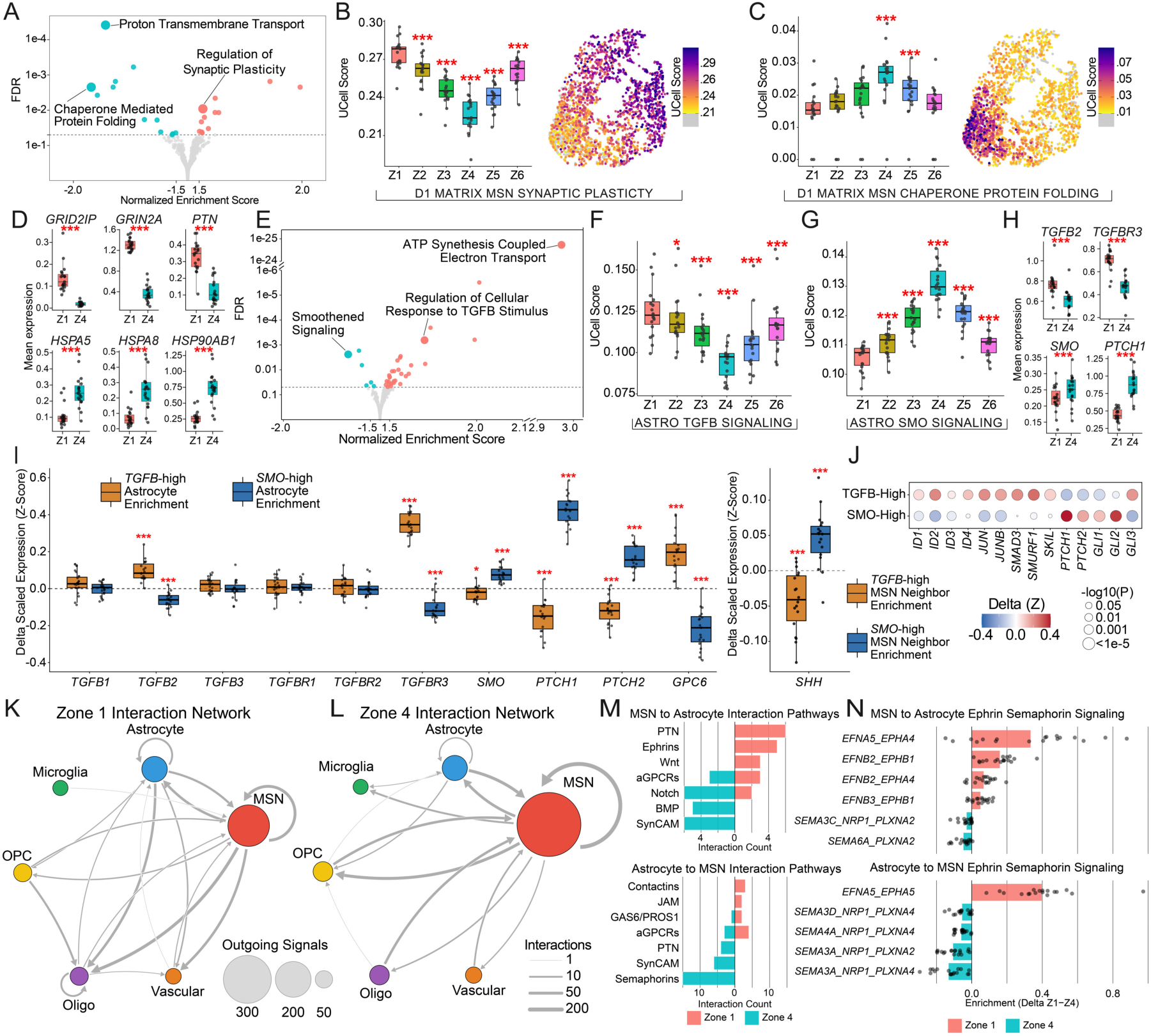
Striatal zones exhibit distinct molecular specializations and spatially coordinated neuron-astrocyte signaling. **A)** Volcano plot of Gene Set Enrichment Analysis (GSEA) results comparing D1 Matrix MSNs in Zone 1 versus Zone 4. Highlighted gene sets denote enrichment for synaptic plasticity in Zone 1 (right) and chaperone-mediated protein folding in Zone 4 (left). **B–C)** UCell module scores across all six zones (left) and corresponding spatial expression maps in a representative donor (right) for **(B)** synaptic plasticity and **(C)** chaperone-mediated protein folding gene sets within D1 Matrix MSNs. **D)** Mean expression levels of representative synaptic plasticity genes (*GRID2IP*, *GRIN2A*, *PTN*) in Zones 1 and 4 across all donors. **E)** GSEA volcano plot for astrocytic populations comparing Zone 1 versus Zone 4, highlighting the selective enrichment of TGF-beta stimulus response in Zone 1 and Smoothened (SMO) signaling in Zone 4. **F-G)** UCell module scores across the six zones quantifying (F) TGF-beta signaling and (G) SMO signaling in astrocytes. **H)** Mean expression levels of representative TGF-beta (*TGFB2*, *TGFBR3*) and SMO pathway (*SMO*, *PTCH1*) genes in astrocytic Zones 1 and 4 across all donors. **I)** Neighborhood analysis depicting the delta scaled expression (Z-score) of TGF-beta and SMO signaling components. Expression is shown for astrocytes (left) and their immediately neighboring MSNs (right), stratified by TGFB-High versus SMO-High astrocytic environments. **J)** Dot plot of downstream transcriptional effectors for TGF-beta and SMO signaling in TGFB-High versus SMO-High astrocytes. Dot size indicates statistical significance, and color represents the delta Z-score. **K–L)** Intercellular interaction networks illustrating the number and directionality of ligand-receptor interactions between MSNs and diverse glial populations in **(K)** Zone 1 and **(L)** Zone 4. Node size reflects the total number of outgoing signals, while edge width represents the pairwise interaction count. **M)** Ranked signaling pathways for MSN-to-astrocyte (top) and astrocyte-to-MSN (bottom) communication, comparing the number of differentially active ligand-receptor pairs between Zone 1 and Zone 4. **N)** Differential enrichment of individual ephrin and semaphorin ligand-receptor pairs driving MSN-to-astrocyte (top) and astrocyte-to-MSN (bottom) signaling in Zone 1 versus Zone 4. *p < 0.05, **p < 0.01, ***p < 0.001.

We examined whether these neuronal specializations were mirrored by the local glial architecture. Intriguingly, we observed that dorsal and ventral astrocytes were enriched for signaling pathways that classically act as opposing morphogens during embryonic telencephalic development. Dorsal astrocytes showed marked upregulation of TGF-beta signaling components, including the ligand *TGFB2* and co-receptor *TGFBR3* **(Figures 6E,F, Table S3**).

Activation of TGF-beta signaling in astrocytes is a known driver of synaptic plasticity in rodents^26,27^, supporting the notion that more dorsal striatum is subject to greater degrees of synaptic remodeling. Conversely, reflecting its ventral developmental origin, ventral astrocytes were enriched for Smoothened (SMO) signaling machinery, and notably, MSN expression of the ventralizing morphogen Sonic Hedgehog (*SHH*) was elevated in this zone (**Figures S6G,H**).

Activation of SMO signaling enhances protein chaperone activity in glia^28^ and promotes neuroprotection^29^, aligning with the proteostatic signature observed in neighboring MSNs within this zone.

To confirm the spatial coordination of these programs, we leveraged the spatial resolution of Slide-tags to perform neighborhood analysis. We defined local cellular neighborhoods (200 µm radius) centered on astrocytes classified as “High” or “Low” for their respective signaling modules. We found that astrocytes within TGFB-High environments were enriched for *TGFB2* and *TGFBR3* (**Figure 6I**). The glypican *GPC6*, known to promote synaptic plasticity^30^, was significantly upregulated in TGFB-high astrocytes (**Figure 6I**). The ligand *SHH* was specifically increased in the MSNs immediately neighboring these astrocytes (**Figure 6I**), and downstream effectors *SMO*, *PTCH1*, and *PTCH2* were all enriched in *SMO-*High astrocytes (**Figure 6J**).

Collectively, these results highlight the spatially segregated neuron-astrocyte signaling loops that define the functional specialization of striatal zones.

To further explore the intercellular signaling within zones, we systematically mapped local (<200 µm) ligand-receptor (LR) interactions between MSN and glial populations in Zones 1 and 4 using the CellChat v2 database^31^. This analysis uncovered 773 differentially active interactions. This signaling landscape was dominated by communication between neurons and astrocytes: MSNs were the primary signal senders (69% of all senders), while astrocytes were the dominant glial partner, accounting for 39% of all glial interactions **(Figure 6K,L)**. This analysis confirmed Astrocyte-to-Astrocyte TGFB2 communication in Zone 1 and MSN-to-Astrocyte SHH communication in Zone 4 **(Figure S6I)**.

We next asked how this astrocyte-MSN crosstalk embodies the distinct states observed in each zone. Analysis of the implicated signaling pathways revealed a striking preservation of embryonic dorsoventral patterning gradients. Zone 1 interactions were dominated by Wnt—a classic dorsalizing morphogen—alongside ephrin signaling, pathways classically associated with continuous synaptic plasticity and remodeling **(Figure 6M,N).** In contrast, the ventral Zone 4 was defined by Notch, BMP, and reciprocal enrichment of semaphorin signaling—reflecting the localized morphogen networks that establish ventral identity during development. Looking beyond astrocytes, these morphogen gradients extended to signaling from MSN to all cell types **(Figure S6J)**. Notch and BMP interactions were nearly exclusive to Zone 4, while MSN-to-glia Wnt signaling was restricted to Zone 1. Interestingly, MSN-to-MSN Wnt signaling was present in both zones, driven by distinct ligands (e.g. *WNT16* dorsally versus *WNT3* ventrally; **Figure S6J**). The ephrin-semaphorin switch observed across zones is notable, as ephrins mediate highly dynamic, contact-dependent synaptic assembly^32,33^ while semaphorins integrate into the extracellular matrix to provide more stable, long-lasting structural restriction^34–36^. Together, our findings reveal spatially distinct MSN-astrocyte signaling across striatal zones, with ephrin-semaphorin interactions among the most prominent, suggesting that zone-dependent intercellular communication — spanning diverse signaling modalities — contributes to the maintenance of striatal organization.

### Reduction in zonal distinctions with age

Aging is a primary risk factor for neurodegeneration, yet its effects are not uniformly distributed across the brain. Within the striatum, macroscale neuroimaging and postmortem studies have long pointed to a dorsal-ventral gradient of vulnerability: the dorsal caudate nucleus exhibits more volumetric atrophy compared to ventral areas^37,38^ and is the primary site of initial degeneration in Huntington’s disease and X-linked dystonia-parkinsonism. Moreover, the putamen acts as a site of selective iron accumulation with age, relative to both the caudate and nucleus accumbens ^39^. These clinical observations suggested that the individual transcriptional zones of the striatum may be differentially affected by aging.

To explore this, we leveraged a population-scale snRNA-seq dataset generated from 131 postmortem donors spanning a wide range of ages (**Figure S7A**). Given the robust spatial gene expression signatures identified in our Slide-tags cohort, we reasoned that we could accurately impute zonal positions for the neurons and astrocytes within this dataset (**Figure 7A**). Because the population-scale snRNA-seq dataset was generated from rough anatomical dissections of the caudate, putamen, and accumbens, we first validated the imputation by examining the relative representation of zones within each sample. As expected, Zone 4 was highly enriched in accumbens dissections, while Zones 5 and 6 were enriched in putamen sections (**Figure S7B**). Most importantly, examination of the imputed snRNA-seq data confirmed highly concordant expression of canonical zonal markers (**Figures 7B, S7C**), establishing our ability to confidently map zonal position onto dissociated, population-scale data.

**Figure 7.**
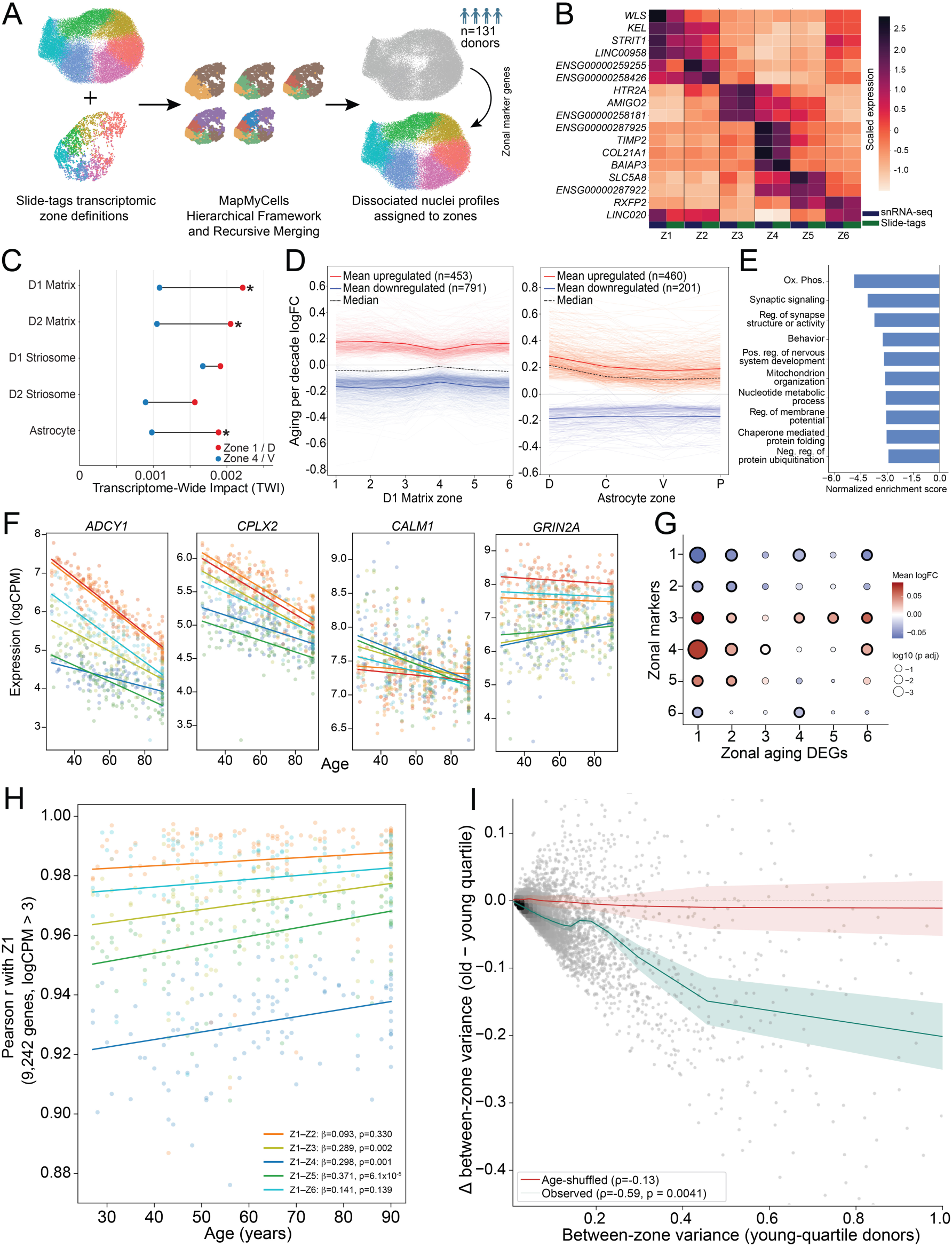
Striatal zones become progressively less molecularly distinct during normal aging. **A)** Schematic of zonal identity imputation onto a population-scale snRNA-seq dataset (n = 131 donors). **B)** Normalized expression of zonal marker genes in the imputed snRNA-seq (blue) and Slide-tags reference (green) datasets. **C)** Transcriptome-wide impact (TWI) for imputed cell types in dorsal (Zone 1/D) versus ventral (Zone 4/V) compartments. Stars indicate significance of dorsal-ventral difference after permutation testing (1,000 iterations, P < 0.002). **D)** Per-decade aging logFC across zones for significantly dysregulated genes (*P*adj < 0.01, |mean logFC| > 0.1) in D1 Matrix MSNs (left, n = 1,244) and astrocytes (right, n = 661). **E)** Gene Ontology (GO) enrichment for the dominant aging expression component identified via ICA (**Figure S7E**). **F)** Mean expression (logCPM) of four exemplar genes across all six zones over the lifespan, with ordinary least squares regression fits. **G)** Enrichment of zonal markers among aging DEGs in D1 Matrix MSNs. Black outlines denote significance (FDR-adjusted *P* < 0.05, two-sided Wilcoxon rank-sum test). **H)** Age-dependent Pearson correlations of average expression (9,242 highly expressed genes, logCPM > 3) between Zone 1 D1 Matrix MSNs and all other zones. **I)** Relationship between baseline cross-zonal expression variance and age-related variance change in D1 Matrix MSNs (n = 9,242 genes). Point shading indicates local gene density. The teal line represents the observed smoothed trend ± 95% CI. The red line denotes the expected null trend ± IQR, derived from permutation testing of young- and old-quartile donors (empirical *P* = 0.0041, 10,000 permutations).

To evaluate zonal differences in aging trajectories, we performed differential expression analysis and calculated an aging TWI for each imputed cell type, within each zone (**Methods**). Within both D1 and D2 Matrix MSNs, a consistent spatial gradient emerged: the lowest aging TWI was observed in the ventral Zone 4, while the highest aging TWIs were localized to the dorsolateral Zone 1 and the dorsal caudate Zone 2 (**Figure 7C**). Interestingly, dorsal astrocytes in Zone 1 mirrored their neuronal neighbors, exhibiting a higher aging-associated transcriptional drift than astrocytes in Zone 4. Striosomal MSNs followed this same trend toward a dorsal bias, though the results did not reach statistical significance. Evaluation of the strongest differentially expressed genes (DEGs) in MSNs revealed that aging was biased toward gene downregulation (64% downregulated, p_adj < 0.01, |logFC| > 0.1, **Figure 7D**), with Zone 4 exhibiting the most compressed aging effects for both up- and downregulated genes. In contrast, astrocytic aging was biased toward gene upregulation (70% upregulated, p_adj < 0.01, |logFC| > 0.1, **Figure 7D**), supporting the widespread observation that astrocytes adopt increasingly reactive states during normal aging^40^.

While many MSN DEGs followed this broad trajectory of dorsally biased dysregulation, the effects were not strictly uniform (**Figure S7D**). To distill these profiles into consistent spatial patterns of age-related gene regulation across the D1 Matrix MSNs, we applied independent component analysis (ICA, **Methods**). The dominant aging signature (Component 1, explaining 76.7% of the variance) captured a globally dysregulated transcriptional program across all zones, though its effects were notably blunted in the ventral Zone 4, perfectly mirroring the overall TWI trend (**Figure S7E**). The genes driving this dominant aging component were heavily enriched for pathways governing cellular metabolism (including complex I and nuclear-encoded mitochondrial genes), protein chaperones, and synaptic plasticity (**Figures 7E, S7F**). The coordinated downregulation of multiple gene sets originally identified as spatial drivers of zonal identity—particularly synaptic plasticity, protein chaperones, and oxidative phosphorylation pathways—suggested a hypothesis: that zonal specialization itself degrades with age. Indeed, several of the top age-downregulated synaptic genes, such as *ADCY1*, *CPLX2*, *CALM1*, and *GRIN2A*, strongly recapitulated this trend, with their highly distinct zonal expression averages progressively converging across the lifespan (**Figure 7F**).

To systematically evaluate this, we examined the enrichment of zonal markers within our aging DEGs. In the dorsal territories (most notably Zone 1), resident markers were globally decreased, while markers of ventral zones tended to aberrantly increase, suggesting a spatial blurring of cellular identity (**Figure 7G**). Furthermore, when correlating the average expression of each zone’s Matrix MSNs against every other zone using the 9,242 most highly expressed genes, cross-zonal correlations systematically increased with age (**Figures 7H, S7G**). This homogenization trend was driven by reduction in between-zone variance for 88.2% of all 27,730 tested genes (**Figure S7H,** permutation p = 0.005, 10,000 iterations). Strikingly, the magnitude of a gene’s baseline spatial variance strongly predicted its vulnerability: the more strongly a gene defined a specific zone, the more aggressively its expression compressed with age (**Figure 7I**). Collectively, these results reveal that aging drives a progressive spatial dedifferentiation of the human striatum, eroding the transcriptional boundaries that separate its specialized functional zones.

## Discussion

Despite its seemingly homogeneous cytoarchitecture, the human striatum spatially segregates into functionally distinct territories. By deploying Slide-tags across a cohort of 19 human donors, we mapped the robust and consistent features of this micro- and mesoscale compartmentalization. Our unbiased spatial sampling uncovered a novel, highly clustered MSN subtype, D1 KCNJ6. Alongside the canonical striosomes and STRv NUDAP populations^41,42^, this discovery adds even further diversification to striatal compartments. This "archipelago" architecture may be driven by computational and neurochemical considerations. Physical aggregation optimizes wiring economy and supports circuit encapsulation^43^, acting as highly specific gating nodes for limbic and autonomic inputs^44,45^. Moreover, tight clustering can create highly concentrated biochemical domains, trapping and pooling local paracrine signals to reach critical activation thresholds^46,47^. Our observation of differential exclusion of interneuron types from striosomal compartments supports the idea that aggregation of rarer, distinct MSN types enables them to establish insulated, distinct circuitry within the broader matrix topography.

Most significantly, our ability to profile the full extent of the human striatum allowed us to uncover a highly conserved mesoscale cytoarchitecture across the human striatum. While the striatum lacks obvious histological boundaries, diverse fields of neuroscience have long identified evidence for spatial territorialization. Anatomical tract-tracing has demonstrated that topographically organized cortical^3–5^, and dopaminergic^48,49^ inputs physically segregate the striatum into parallel domains. In living humans, resting-state functional imaging has similarly defined distinct connectivity networks that parcellate the structure^50^. Intriguingly, the physical boundaries of our zones bear topographical similarities to parcellations defined by connectivity and functional imaging. Broadly speaking, our Zones 2 and 3 correspond to the associative striatum in caudate; Zone 4 corresponds to the region of high limbic input in the accumbens, and Zones 5 and 6 overlap with the sensorimotor region of the putamen. Zone 1–which spans the dorsal tips of both the caudate and putamen–overlaps regions annotated as both sensorimotor and associative. Ultimately, mapping these intrinsic cellular microenvironments onto the primate connectome will provide a definitive, multi-scale understanding of striatal functional neuroanatomy.

One piece of evidence linking our molecularly defined zones to functional networks derives from a shared vulnerability to aging. We found that the spatial boundaries defining Matrix MSN zones progressively “blur” over the human lifespan, as MSNs lose their highly specialized signatures and converge toward a more homogeneous transcriptional state. Intriguingly, functional MRI studies have observed an analogous macroscopic pattern in the aging striatum, where older adults exhibit reduced decoupling of the caudate from the default mode network^51^, decreased specificity in functional coupling between the caudate and cortical association networks^52^, and a reduction of functional specialization for implicit learning tasks^53^. This so-called “dedifferentiation” in functional imaging mirrors our observations, suggesting that these molecular and functional phenomena may be intrinsically linked.

A particularly intriguing feature of our analysis of the molecular pathways underlying striatal zones was the discovery of persistent developmental morphogen gradients acting as spatial organizers in the mature human brain. During embryogenesis, the specification of the dorsal-ventral axis in the developing forebrain is tightly governed by orthogonal, competing morphogen signals: the dorsal pallium is patterned by Wnt and TGF-beta gradients, while the ventral telencephalon is specified by Hedgehog and Notch signaling^54^. Our spatial atlas indicates that this primordial biochemical architecture persists as an actively maintained homeostatic scaffold in the adult human striatum, and that it erodes with aging. The maintenance of developmental gradients in the adult brain–including Hedgehog^55,56^, Notch^57,58^, and Wnt^59^–has been reported in the mouse, while broader embryonic gradients have been observed in bulk RNA-seq data gathered across the human brain^60^. Crucially, this provides a compelling mechanistic hypothesis for why our zonal boundaries progressively ‘blur’ over the lifespan: the spatial dedifferentiation of the aging striatum may fundamentally represent the progressive erosion of the brain’s lifelong morphogenic scaffolding, leading to a collapse of regional functional specialization.

Beyond the specific biology of the striatum, our work establishes an experimental and analytical framework for constructing a robust, data-driven neuroanatomy. Two specific components of our strategy were crucial to resolving striatal zones. First, the transcriptome-wide, single-nucleus profiling afforded by Slide-tags allowed us to detect a subtle topology in the gene expression latent space–namely, the “ring-like” architecture of MSNs–that would be obscured by signal mixing inherent to other spatial technologies. Second, by profiling a cohort of 19 donors, we replaced somewhat arbitrary clustering parameters (often defined by the number of differentially expressed genes between clusters, for example) with an empirical standard: the maximum granularity that preserves spatial consistency across individuals. This criterion redefines “cell types” and “territories” not as subjective computational choices, but as reproducible biological features conserved across the human population. We envision this framework as the foundation for the construction of a complete, quantitative, and spatially resolved neuroanatomy of the human brain that has previously been out of reach.

### Limitations of the study

To confidently define and spatially position robust, cross-donor reproducible cell types and states, our spatial profiling focused primarily on the anterior striatum. Although our initial profiling of the human posterior striatum and the macaque indicates that zonal definitions are preserved across the structure and across primate evolution, a broader sampling of the basal ganglia, and expansion across a wider phylogenetic cohort, will be required to definitively map all zonal boundaries. Additionally, to secure a sample size necessary to model lifespan aging trajectories, we utilized computational imputation to map zonal identities of the most molecularly distinct populations (MSNs, astrocytes) onto a population-scale snRNA-seq cohort. Further improvements to Slide-tags throughput will enable case-control spatial genomics studies to reach sample sizes that will allow direct measurement, bypassing the need for imputation and enabling the physical mapping of age-related transcriptomic drift and disease-associated vulnerabilities across all cell types.

## Acknowledgements

We thank Gord Fishell and Bernardo Sabatini for thoughtful comments on our manuscript. This work was supported by the National Institutes of Health/National Institute of Mental Health (BRAIN UM1MH130966 to S.A.M. and E.Z.M), Alzheimer’s Association (AACSF-1241991 to A.W.K), the Mass General Brigham Neuroscience Institute (A.W.K), the Heitman Foundation (A.W.K), and the Stanley Center for Psychiatric Research.

## Data and Code Availability

A viewer of the Slide-tags data–to investigate average and spatial patterns of gene expression variation across zones–is available at https://storage.googleapis.com/capalpha/index.html.

Raw data, including fastq files, counts matrices, and spatial placements are available through the NIH Brain Initiative’s Neuroscience Multiomic Data Portal: https://assets.nemoarchive.org/collection/nemo:dat-aqicdbo.

The Slide-tags computational workflow to spatially place profiled cells is available at: https://broadinstitute.github.io/warp/docs/Pipelines/SlideTags_Pipeline/README.

## Author Contributions

E.Z.M., K.I., and S.A.M. conceived of the study. A.W.K. and M.L. performed the analyses, with help from M.S., E.M., O.Y., K.S., L.R., J.N., S.B., and E.Z.M. N.A.R. led the generation of the Slide-tags data, along with J.M., H.G., and C.V., and with help from K.B., N.N., O.C., A.M., and N.B. V.K. synthesized the barcoded Slide-tags beads. K.F. and E.F. assisted with study design and implementation. A.W.K. and E.Z.M. wrote the manuscript, with input from all authors.

## Declaration of Generative AI and AI-assisted Technologies

During the preparation of this work, the authors used Claude, Gemini, and ChatGPT for building syntactically correct code, debugging errors, and improving computational efficiency. Gemini was also used for editing and proofreading of manuscript text. After using these tools, the authors reviewed and edited the content and take full responsibility for their outputs.

---

**Figure S1.**
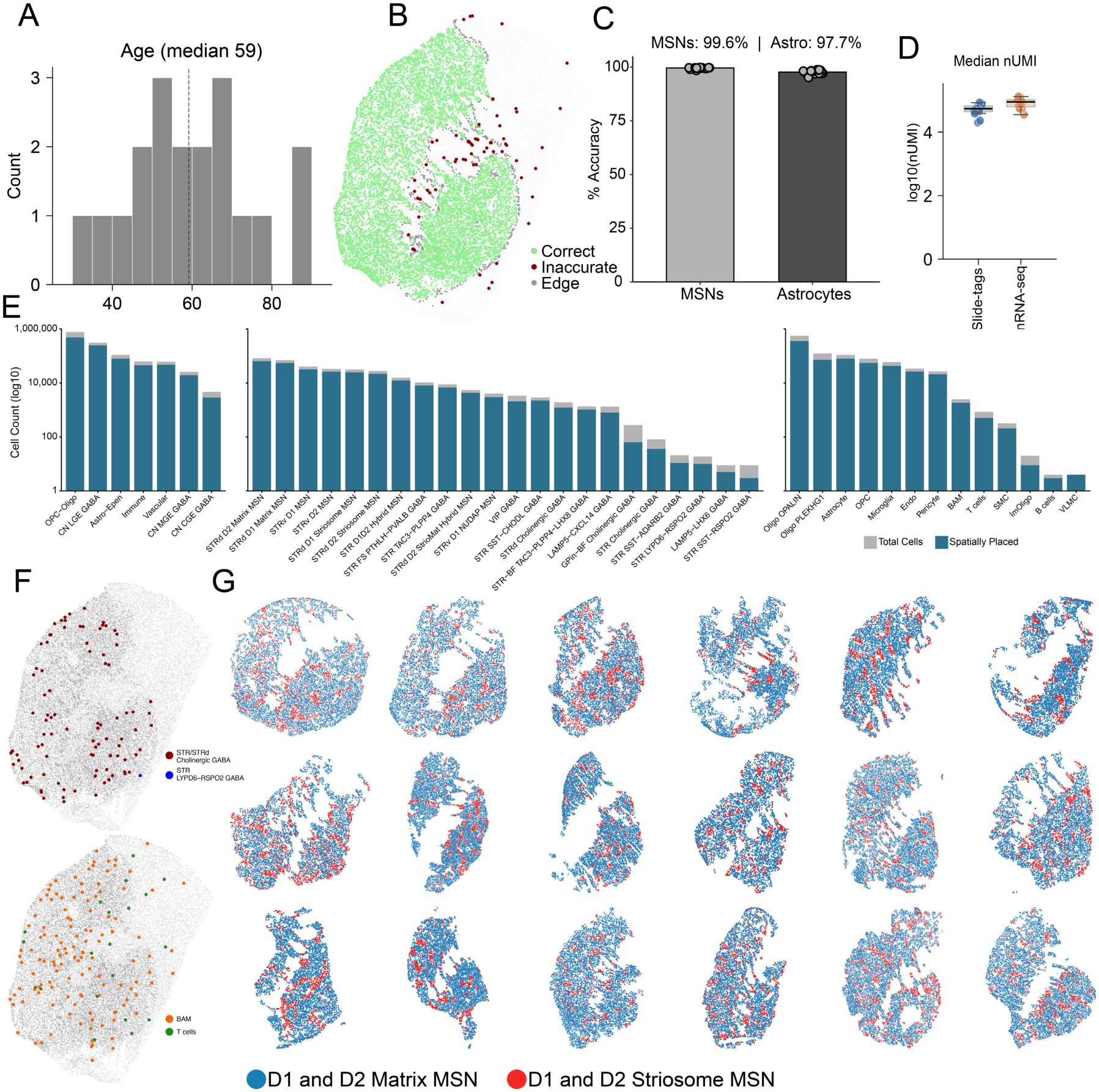
Cohort demographics, technical validation, and spatial cellular census of the Slide-tags human striatal atlas. **A)** Age distribution of the 19 postmortem donors profiled within the spatial cohort. **B)** Spatial mapping specificity evaluated in a representative donor. The medium spiny neuron (MSN) false-placement rate was quantified utilizing white matter tracts as an anatomical negative control (profiles inaccurately mapping to white matter are flagged in red). **C)** Quantification of spatial mapping accuracy for MSNs and astrocytes across the full 19-donor cohort, demonstrating highly specific spatial localization. **D)** Transcript capture sensitivity, displayed as the median unique molecular identifiers (nUMI) per MSN. Transcript recovery in the Slide-tags spatial cohort (n=19 donors) is comparable to a reference standard single-nucleus RNA-seq cohort (n=131 donors). **E)** Yield of total recovered and spatially localized nuclei across major cell classes (left), fine-grained neuronal subtypes (middle), and non-neuronal subtypes (right), annotated using a reference basal ganglia taxonomy^18^. **F)** Representative spatial localizations of selected neuronal (top) and non-neuronal (bottom) populations. **G)** Spatial distribution of the cardinal Matrix and Striosome MSN compartments across the remaining 18 donors. **H)** Inhomogeneous pair correlation (g_inhom(r)) evaluated against a background-fixed simulation envelope. Simulation envelopes (gray ribbons) were generated from 199 Monte Carlo simulations of an inhomogeneous Poisson process under a background density estimated from all MSN cells; exceedance of the envelope indicates significant spatial clustering beyond tissue-level MSN density gradients.

**Figure S2.**
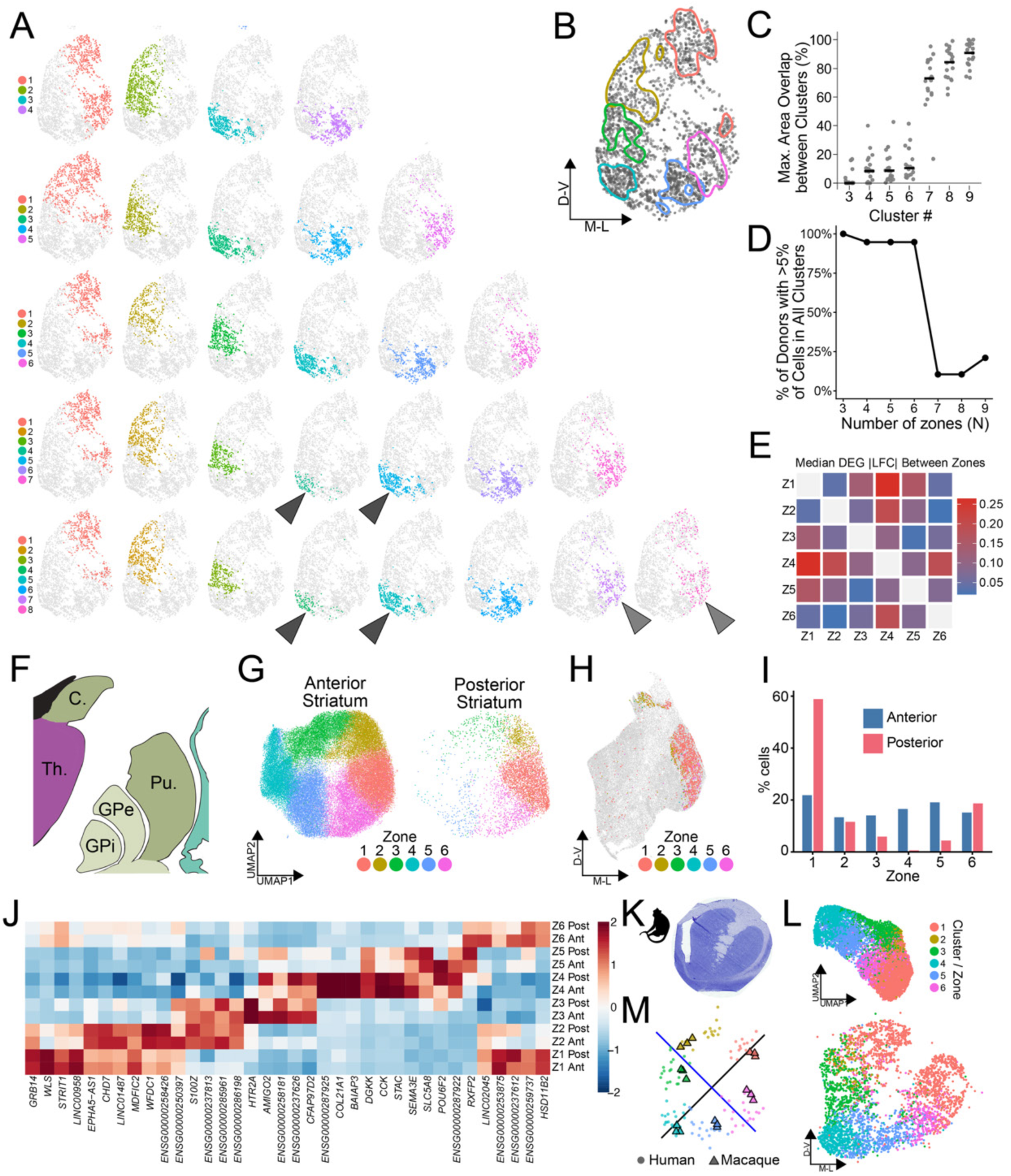
Additional spatial and cross-species validation of D1 Matrix zones. **A)** Spatial distribution of *k*=4 through *k*=8 clustering solutions in a representative donor (arrowheads: examples of high spatial overlap). **B)** KDE-derived contour maps of zonal territories. **C)** Maximum spatial overlap area between cluster pairs for each *k* solution. **D)** Minimum donor representation within any cluster as a function of cluster count**. E)** Median log-fold change magnitude of differentially expressed genes across all zone pairs. **F)** Anatomical diagram of the posterior striatum region profiled by Slide-tags (adapted from Ding et al., 2016^61^). **G)** Joint UMAP embedding of anterior (19 donors, left) and posterior (2 donors, right) D1 Matrix MSNs. **H)** Spatial distribution of posterior D1 Matrix MSNs in a representative donor. **I)** Percentage of D1 Matrix MSNs assigned to each zone in anterior versus posterior sections. **J)** Normalized expression of D1 Matrix zonal markers across anterior and posterior sections. **K)** Nissl stain of a coronal *Macaca fascicularis* striatum section adjacent to the Slide-tags assay. **L)** UMAP embedding (top) and corresponding spatial map (bottom) of macaque D1 Matrix neurons. **M)** Aligned spatial centroids for each zone across the 19 human donors (circles) and 3 adjacent macaque sections (triangles).

**Figure S3.**
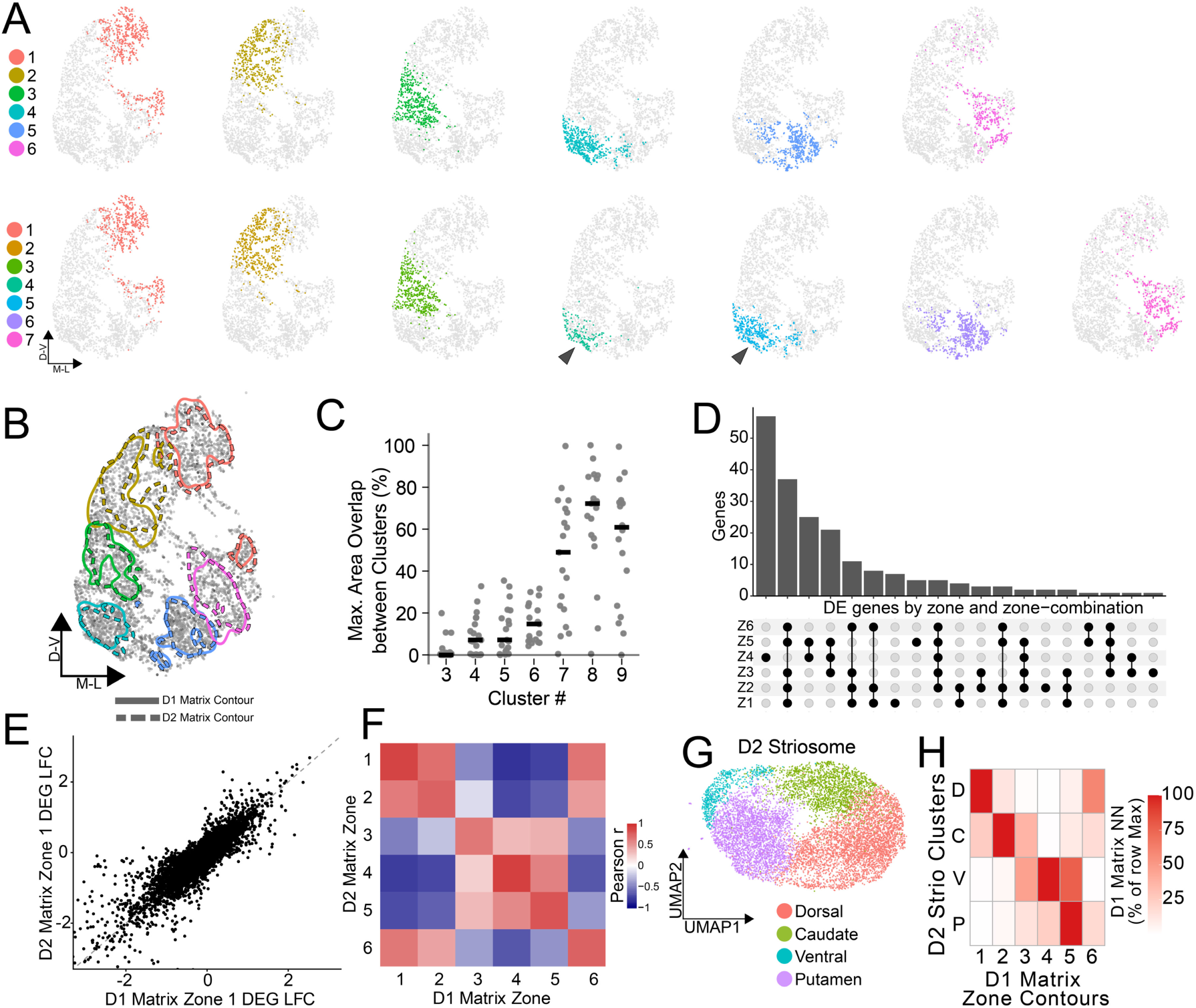
Extended spatial analysis of diverse MSN subtypes. **A)** Spatial distribution of *k*=6 (top) versus *k*=7 (bottom) clustering solutions for D2 Matrix MSNs in a representative donor. **B)** Contour maps comparing boundaries defined by D1 versus D2 Matrix MSNs. **C)** Maximum spatial overlap area between D2 Matrix cluster pairs across *k* solutions. **D)** UpSet plot detailing differentially expressed genes (DEGs) shared across single or combined D2 Matrix zones. **E)** Scatter plot comparing log-fold changes of Zone 1 marker genes between D1 and D2 Matrix MSNs. **F)** Cross-population correlation of zonal markers between D1 and D2 Matrix MSNs. **G)** UMAP embedding of D2 Striosome MSNs. **H)** Spatial contour overlap between D2 Striosome clusters (rows) and D1 Matrix zones (columns).

**Figure S4.**
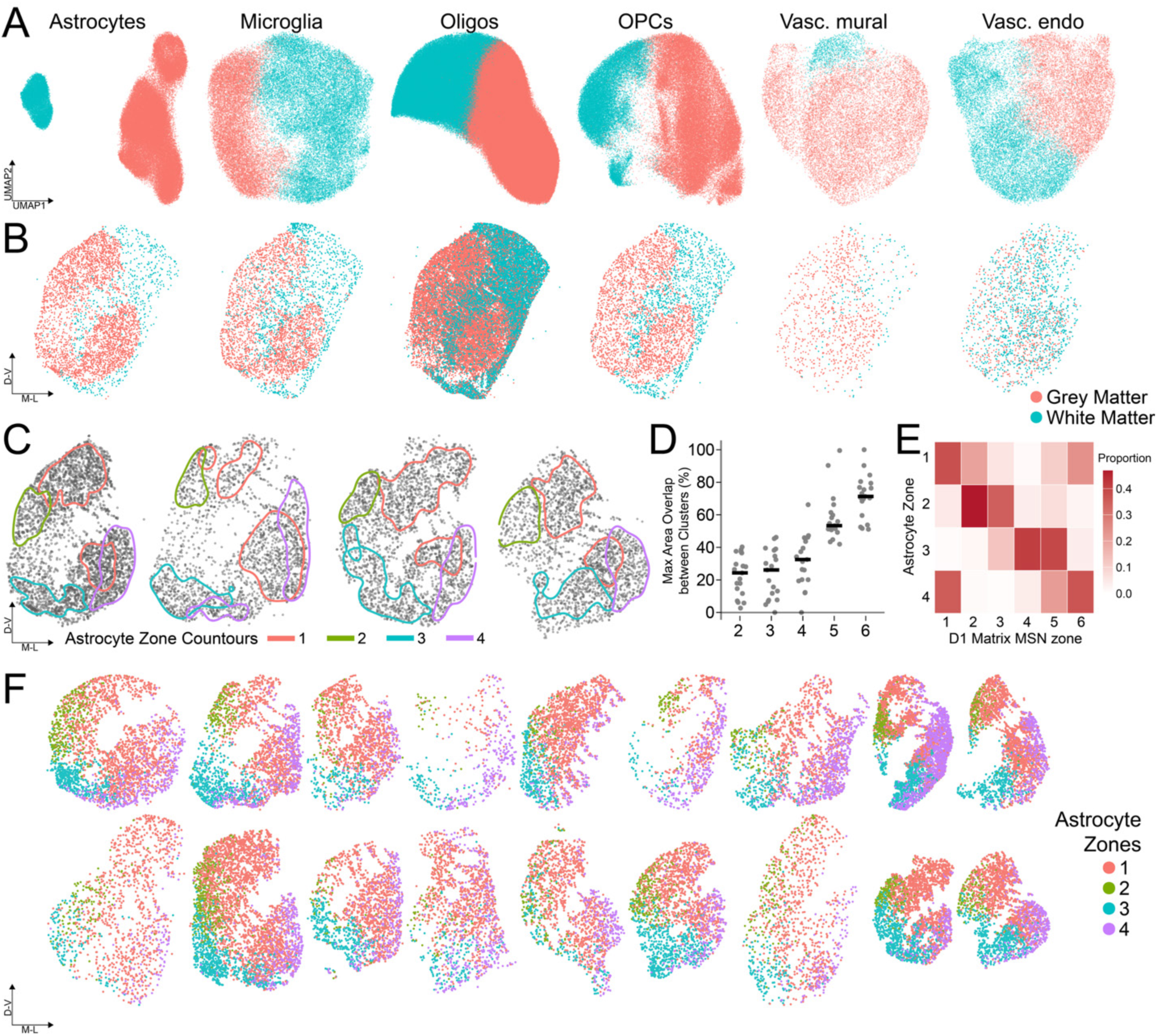
Spatial parcellation of striatal glia. **A)** UMAP embeddings and spatial **(B)** maps of glial populations distinguishing gray versus white matter identity. **C)** Contour maps of the four astrocyte zones identified via unsupervised clustering. **D)** Maximum spatial overlap area between astrocyte cluster pairs across *k* solutions. **E)** Spatial contour overlap between astrocyte clusters (rows) and D1 Matrix MSN zones (columns). **F)** Spatial distribution of zonated astrocytes across all 19 donors.

**Figure S5.**
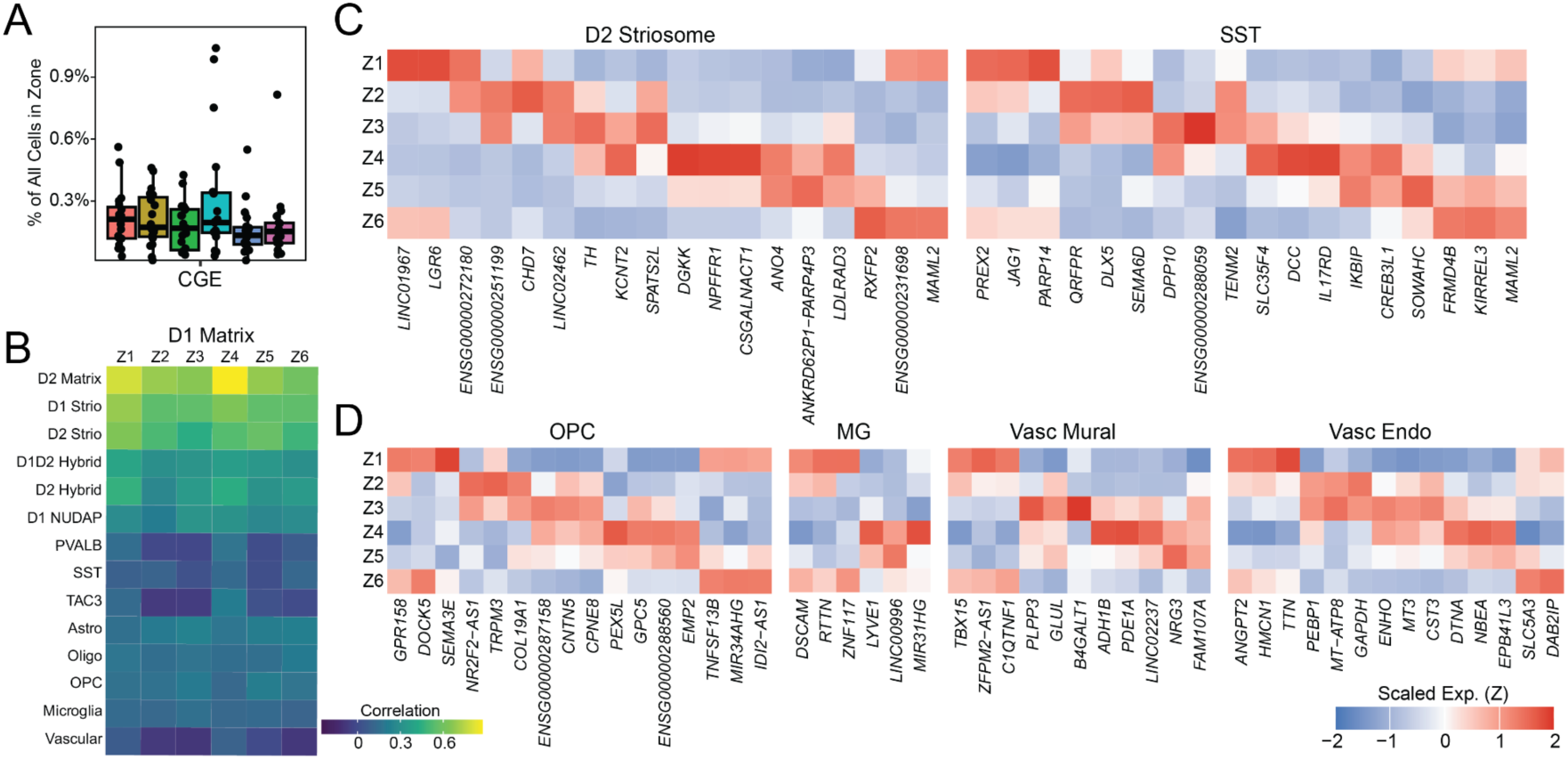
Striatal zones possess distinct cellular architectures and transcriptional specializations. **A)** CN CGE GABA neuron proportions for each sNN assigned zone. **B)** Cross-cell-type correlation of expression changes (log-fold change) for the union of zonal marker genes, comparing D1 Matrix MSNs to all other cell populations. **C-D)** Heatmap of normalized expression for representative zone-specific marker genes for **(C)** D2 Striosome MSNs, SST interneurons and **(D)** glial cells within each D1 Matrix-defined striatal zone.

**Figure S6.**
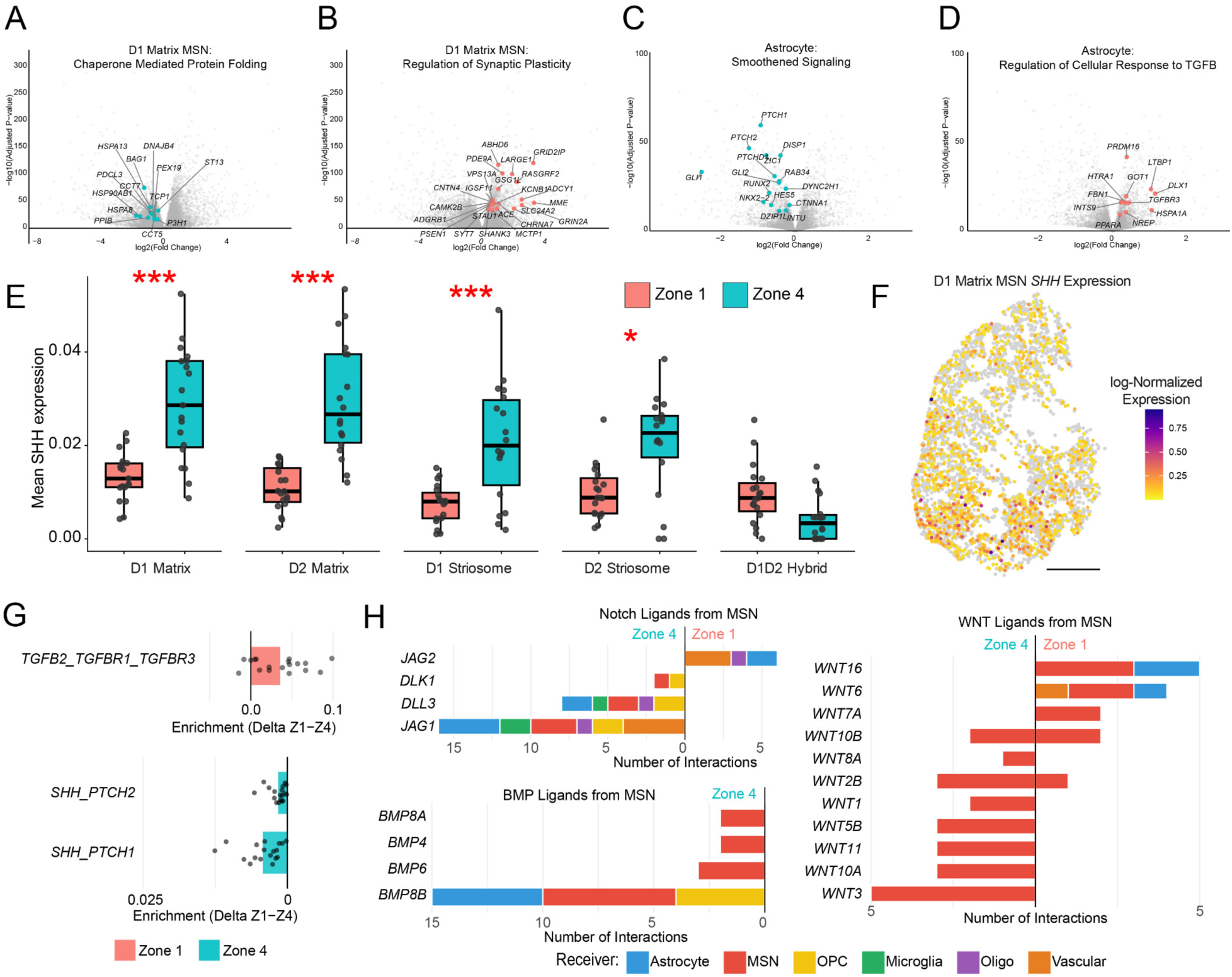
Zone-specific gene set enrichment in D1 Matrix MSNs and astrocytes. **A,B)** Volcano plots of DESeq2 results highlighting the top 30% of leading-edge genes from **(A)** Chaperone-Mediated Protein Folding and **(B)** Regulation of Synaptic Plasticity. **C,D)** Volcano plots of DESeq2 results highlighting the top 30% of leading-edge genes from **(C)** Smoothened Signaling and **(D)** Regulation of Cellular Response to TGF-β. **(E)** Mean expression levels of MSN *SHH* in Zones 1 and 4 across all donors (*p < 0.05, **p < 0.01, ***p < 0.001), and **(F)** corresponding spatial D1 Matrix MSN *SHH* expression map in a representative donor (scale bar = 0.5cm). **G)** Differential enrichment of *TGFB2* ligand interaction with the *TGFBR1* and *TGFBR3* complex in Astrocyte-to-Astrocyte signaling (top) and *SHH* ligand interactions in MSN-to-Astrocyte signaling (bottom). **H)** Interactions ranked by ligand for MSN-to-any cell type for Notch signaling (top left) and BMP signaling (bottom left) and Wnt (right) comparing the number of interactions between Zone 1 and Zone 4.

**Figure S7.**
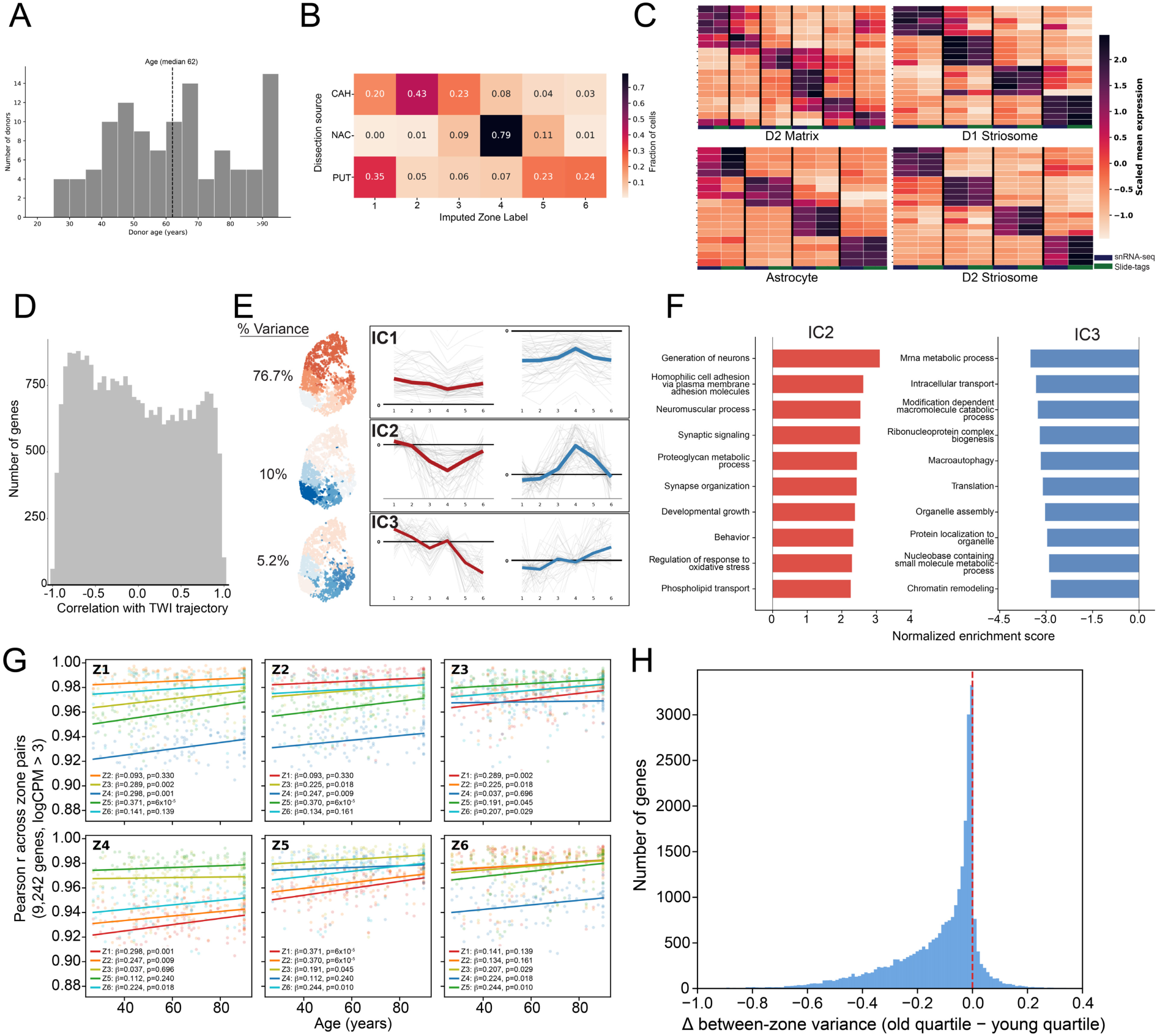
Validation of zonal imputation and expanded spatial aging analyses. **A)** Age distribution of the 131-donor snRNA-seq cohort. **B)** Confusion matrix comparing the macroscopic anatomical dissection region to the computationally imputed spatial zone. **C)** Normalized expression of canonical zonal markers across imputed D2 Matrix, D1 Striosome, D2 Striosome, and Astrocyte populations. **D)** Distribution of correlation coefficients between aging DEG logFC values and TWI across zones for all MSNs. **E)** Independent component analysis (ICA) of aging DEGs in D1 Matrix MSNs. Plots display the individual and average component loadings for the 50 highest- (red) and lowest-scoring (blue) genes. **F)** Gene Ontology (GO) pathway enrichment for Component 2 (left) and Component 3 (right). **G)** Age-dependent Pearson correlations of average gene expression (9,242 highly expressed genes, logCPM > 3) between all pairwise spatial zone combinations. **H)** Distribution of the change in between-zone expression variance (Δ variance) for D1 Matrix MSNs, comparing the oldest quartile (n = 30 donors, age > 75) to the youngest quartile (n = 28 donors, age < 49). 88.2% of genes (n=27,730) exhibit a negative Δ variance, indicating global, age-driven spatial compression. Statistical significance (P = 0.005) was assessed via 10,000 permutations shuffling donor age labels.

## Supplemental tables

**Table S1: Donor demographics and sequencing metrics for Slide-tags experiments.** Age, sex, total cell count, median nUMI across all cells, and median nUMI for medium spiny neurons (MSNs) for each donor used in this study.

**Table S2: Gene set enrichment analysis results for Zone 1 vs. Zone 4 D1 matrix MSNs.** Full GSEA results for GO Biological Process gene sets comparing Zone 1 and Zone 4 D1 Matrix MSNs. The workbook contains three sheets: Kept_terms, significant gene sets retained after redundancy pruning; All_sig_terms, all significant gene sets with pruning status and the representative pathway each pruned term was grouped with; and All_terms, all tested gene sets. Leading-edge genes are listed for each term.

**Table S3: Gene set enrichment analysis results for Dorsal vs. Ventral astrocytes.** Full GSEA results for GO Biological Process gene sets comparing Dorsal and Ventral Astrocytes. The workbook contains three sheets: Kept_terms, significant gene sets retained after redundancy pruning; All_sig_terms, all significant gene sets with pruning status and the representative pathway each pruned term was grouped with; and All_terms, all tested gene sets. Leading-edge genes are listed for each term.

**Table S4: Spatially enriched ligand-receptor interactions.** Significant and robust ligand-receptor interactions identified through zonal interaction analysis, comparing zone 1 versus zone 4.

## Methods

### Donor selection and tissue procurement

Postmortem human brain tissue was obtained through the Human Brain Cell Atlas Collection (HBCAC), a program within the NIH NeuroBioBank (NBB) supporting the BRAIN Initiative Cell Atlas Network (BICAN). Contributing repositories followed harmonized protocols for donor evaluation and tissue processing.

Tissue collection adhered to the United States Uniform Anatomical Gift Act and applicable laws, with informed consent for unrestricted research use obtained from the legal next-of-kin by the NBB. The Broad Institute Office of Research Subject Protection determined that the use of these de-identified specimens does not constitute human subjects research requiring IRB review (NHSR-8066). Adult donors (ages 26 to 90+, Table S1) were systematically evaluated and excluded if they possessed major neurological or psychiatric conditions (e.g., Alzheimer’s disease, schizophrenia) or positive serology for HIV-1/2, Hepatitis B, or C. Donors with minor, non-debilitating conditions (e.g., migraines), historical diagnoses in clinical remission lacking confirmatory neuropathology, or incidental focal pathology associated with normal aging were permitted.

To ensure standardized spatial sampling, scaled photographs of 5-mm coronal brain slabs were evaluated using the Neuroanatomy-anchored Information Management Platform (NIMP, UTHealth Houston). The striatum target region, located approximately 5.5 cm from the anterior pole on the 6th coronal slab, included the head of the caudate, the putamen anterior to the anterior commissure, and the nucleus accumbens^61^. Expert neuroanatomists identified slabs containing the relevant striatal regions and digitally annotated precise regions of interest (ROIs) linked to the Harmonized Ontology of Mammalian Brain Anatomy (HOMBA). To enforce experimental uniformity, ROIs were strictly matched across all donors, minimizing sampling variability introduced by individual macroscopic morphological differences.

### Barcoded bead array fabrication and reconstruction

Bead synthesis and array fabrication were performed as previously described^16,17^ with the following modifications: spatial DNA barcodes were extended to 17 total bases. The capturing and releasing bead sequences were:

Capture beads: 5′-TTTTCTACACGACGCTCTTCCGATCTJJJJJJJJJTCTTCAGCGTTCCCGAGAJJJJJJJJNNNNNNNNNTTT CTTATATGGG-3′

Spatial barcode release beads: 5′-TTT-pc-GTGACTGGAGTTCAGACGTGTGCTCTTCCGATCTJJJJJJJJJJJJJJJJJCTGTTTCCTGNNNNNNNNNCC CATATAAGAAA-3′

To fabricate the arrays, round glass coverslips were cleaned and functionalized by immersion in charging buffer (8:2 ratio of 200-proof ethanol to 10% glacial acetic acid, supplemented with 0.2% v/v 3-(trimethoxysilyl)propyl methacrylate [Sigma, 440159]) for ≥10 min. Coverslips were subsequently rinsed three times in 200-proof ethanol and dried. To form the array substrate, a 4% acrylamide gel solution (40% Acrylamide/Bis-acrylamide 19:1 [Bio-Rad 1610144], 0.06% v/v 10% ammonium persulfate, and 0.06% v/v TEMED in ultrapure H₂O) was sandwiched between a functionalized and a non-treated coverslip. Following a 40–60 min polymerization at room temperature, coverslips were separated, and any arrays lacking uniform gel coverage were discarded.

Donut-shaped silicone gaskets (1.6 mm non-adhesive silicone sheet; inner diameter 30 mm; Sigma, GBL664273) were then affixed to the gel surface. Capture and releasing beads were pooled at a 1:3 ratio and resuspended in 10% DMSO at a concentration of 16,000–28,000 beads/µL. A 900 µL volume of this suspension was applied to the gasketed array and covered with a thin, non-adhesive silicone sheet to eliminate air bubbles. After a 3 min settling period, the array was gently tapped and reinoculated to ensure uniform monolayer formation. Finally, the silicone sheet and gasket were removed, excess beads were washed away with indirect ultrapure water, and the arrays were air-dried.

To generate a sequencing library for the computational reconstruction of physical bead coordinates, arrays underwent a controlled diffusion process as previously described^17^. Arrays were washed and subsequently submerged in 800–1000 µL of 2× SSC buffer (Invitrogen, 15557-004). Oligonucleotides were cleaved from the releasing beads via a 12-s exposure using a PX2 365 nm LED photoreactor (Alpha Thera). The arrays were then incubated in the dark for 10 min to permit the released nucleotides to crosslink with adjacent capture beads. Following a 2× SSC wash, arrays were incubated in an extension buffer (1× NEBuffer 2 [NEB, B7002S], 1 mM dNTPs [NEB, N0447L], and 0.15 U/µL Klenow Exo⁻ [NEB, M0212L]) at 37 °C for ≥1 h. Arrays were washed twice more with 2× SSC, submerged in 1 mL of ultrapure H₂O, and heated to 95 °C for 5 min on a dry block to elute the extended barcodes. The resulting eluate was immediately collected.

Eluted melt products were amplified using 1× KAPA HiFi polymerase (Roche, KK2612) with 100 nM P5-Truseq Read 1 and 100 nM P7-Truseq Read 2 primers. PCR cycling conditions were: 95 °C for 3 min; 4–9 cycles (titrated based on estimated barcode yield) of 98 °C for 20 s, 65 °C for 15 s, and 72 °C for 15 s; followed by a final extension at 72 °C for 1 min and a 4 °C hold. The final spatial barcode libraries were purified using 1.5× SPRISelect beads (Beckman Coulter, A63881).

### Slide-tags procedure

Slide-tags was performed largely as described previously^16^. Fresh-frozen tissue blocks were mounted in OCT and sectioned at 20 µm on a cryostat (Leica CM1950) maintained between −15°C and −25°C.

Sections were transferred directly onto the prepared Slide-tags arrays, immediately hydrated with ice-cold PBS (Thermo Fisher, 10010023), and placed on a pre-chilled (−20°C) metal block. To release the spatial barcodes, the array and cold block were centered under a 365 nm UV light source (ThorLabs, M365LP1-C5) for 2 min. The array was then incubated on the cold block for an additional 7 min to allow the cleaved barcodes to diffuse into the adjacent tissue nuclei.

The array was then transferred to a pre-chilled petri dish, and the tissue was dissociated from the surface via manual trituration in Buffer A (Minute Single Nucleus Isolation Kit; Invent Biotechnologies, BN-020).

Nuclei were pelleted (500 × *g*, 5 min) and the supernatant was discarded. The resuspended pellet was incubated at −20°C for 10 min, passed through a filter tube (13,000 × *g*, 30 s), and the flow-through was centrifuged again (600 × *g*, 5 min). Isolated nuclei were resuspended in PBSAi (PBS supplemented with 5% BSA and 1 U/µL RNase inhibitor) and overlaid onto 1 mL of Buffer B (Invent Biotechnologies, BN-020). Following a final density-gradient centrifugation (1,000 × *g*, 10 min), the concentrated nuclei were stained with 1:1000 DAPI (Thermo Fisher, 62248) in PBSAi and quantified using a Fuchs-Rosenthal disposable hemocytometer (INCYTO, DHC-F01-5) on a fluorescence microscope (Zeiss Axio Observer) prior to 10x GEM generation and library construction.

To prepare Slide-tags nuclei for sequencing, 37.5 µL of nuclei suspension was loaded onto a Chromium Controller X (10x Genomics) using the GEM-X Single Cell 5′ Reagent Kit v3 with Feature Barcode technology for Cell Surface Protein. Procedures strictly followed the manufacturer’s protocol (CG000734, Rev A) with the following modifications for spatial barcode recovery. To specifically capture the Slide-tags spatial barcodes, 1 µL of a custom primer (5′-[GTGACTGGAGTTCAGACGT]-3′) was spiked into the standard cDNA amplification reaction. The cDNA was subsequently amplified for 13-15 PCR cycles.

GEX index PCR was performed for the 5′ gene expression libraries according to the manufacturer’s protocol. Spatial barcode libraries were generated following the 10x cell surface protein library protocol with 12 PCR cycles. To minimize gene expression fragment carryover and specifically enrich for the shorter spatial barcode fragments, a custom double-sided SPRISelect (Beckman Coulter) size selection (0.6× and 1.2× volumetric ratios) was applied to the final barcode libraries prior to sequencing. Gene expression and spatial barcode libraries were sequenced on the Illumina NovaSeq X, 25B flow cell. Gene expression and spatial barcode libraries were sequenced with 28 × 90 bp paired-end reads.

### Histological staining

Following collection of tissue for Slide-tags, an adjacent 10 or 20 µm section was cut for Nissl staining to provide an anatomical reference for spatial transcriptomic data. Sections were equilibrated to room temperature, fixed in 70% ethanol for 2 minutes, and rehydrated in ultrapure water for 30 seconds.

Excess water was removed and slides were stained with freshly filtered 0.1% Cresyl Violet acetate (Electron Microscopy Sciences, 26089-01) for 3–4 minutes. Excess stain was removed and slides were rinsed in ultrapure water for 30 seconds, then sequentially dehydrated in 70%, 95%, and 100% ethanol for 30 s, 30 s, and 1 min, respectively. Slides were cleared in xylene for 5 minutes, mounted with Permount (Fisher Chemical, SP15-100), and coverslipped. Images were acquired using the Keyence BZ-X810 microscope under brightfield illumination at 4× and 10× magnification.

### Sequence alignment

Human sequence data were aligned to the GRCh38 reference genome with GENCODE v43 gene models, and macaque (*Macaca fascicularis*) sequence data were aligned to the Macaca_fascicularis_6.0 genome assembly (GCA_011100615.1; Baylor College of Medicine) with Ensembl gene annotation (release 104), both following the standard Drop-Seq protocol ^62^. Expression matrices were quantified using the Drop-Seq DigitalExpression program with parameters configured to replicate STARsolo quantification ^63^. Ambient RNA was removed from gene expression matrices using CellBender (v0.3.2) remove-background. Nucleus-containing droplets were distinguished from empty droplets using DropSift^64^.

### Spatial array computational reconstruction

The spatial bead coordinates were computationally reconstructed from the sequencing library output using a workflow of scripts written in Julia 1.10.5 and Python 3.11.2. The first script parses through the FASTQ sequences, discards low-quality reads, aggregates reads into UMIs, and calls valid bead barcodes. The result is a spatial barcode count matrix which records the number of observed UMIs between the releasing and capturing bead barcodes. This matrix is converted into a k-nearest neighbor (kNN) graph by evaluating the cosine similarity between each pair of releasing beads and sparsifying by subsetting to the top 150 closest neighbors. To boost the speed and accuracy of the reconstruction process, sensible starting locations for each bead are estimated by partitioning the kNN graph into leiden clusters and running them through a quick UMAP (default parameters) to generate an initial embedding. The kNN graph and these initialized positions are then passed into a lengthier UMAP (4x default epochs, 2x default positive and negative samples, all other parameters default) to generate a complete 2D embedding. This embedding is scaled into micrometers and then saved as the final array coordinates.

### Slide-tags data processing and spatial positioning of nuclei

Slide-tags sequencing data were processed using the Slide-tags Pipeline (RRID:SCR_027567^65^, a cloud-optimized workflow implemented in WDL and executed on the Terra platform. The pipeline integrates three subworkflows. The Optimus workflow aligned gene expression reads to the reference genome using STARsolo and produced cell-by-gene count matrices. The SpatialCount workflow processed spatial barcoding reads to extract spatial barcodes, perform barcode error correction, and map reads to spatial bead coordinates. The Positioning workflow integrated gene expression and spatial data by assigning cell barcodes to spatial bead coordinates, producing a Seurat object with spatial coordinates for downstream analysis.

### HMBA taxonomy assignment of Slide-tags data with MapMyCells

Cell-type labels were transferred from the Human Molecular Brain Atlas (HMBA) basal ganglia taxonomy^18^ using the MapMyCells computational framework (Daniel, Lee, Mollenkopf et al. co-submitted).

### Integrated clustering and quality control

Returned cells were processed using Seurat v5 in R. For each dataset, we calculated quality metrics including number of unique genes (nGene), total unique molecular identifiers (nUMI), percentage of mitochondrial reads (MT%), and percentage of ribosomal subunit reads (Ribo%). Integrated unsupervised clustering was performed on each cell class individually across donors. The top 2,000 variable genes were selected using the variance-stabilizing transformation (vst) method in Seurat. To remove donor-specific batch effects (e.g., due to pre- and postmortem effects, sample preparation, and sequencing), transcriptome data were scaled per gene and per donor (mean of zero and unit variance) prior to principal component analysis. The top 30 principal components, weighted by variance explained, were used for all analyses. Harmony v1.0^66^ was applied to remove residual batch effects from principal components, with donor specified as the batch variable using default theta and lambda parameters. Batch-corrected components were used for shared nearest neighbor (sNN) graph construction and Leiden clustering at resolutions ranging from 0.4 to 0.8.

### Doublet and low-quality cluster removal

Clusters defined by low-quality metrics (low nGene, low nUMI, high MT%, high Ribo%, or high lncRNA%) were removed. Marker genes for each cluster were identified using Seurat’s FindMarkers function.

Doublet clusters were identified by co-expression of canonical marker genes from distinct cell types and removed. Clustering was repeated iteratively using the same methods to ensure complete doublet removal. These doublet-filtered data were used for all cell proportion and compartment analyses.

### Additional expression quality filtering

For gene expression analyses, additional quality filtering was performed within each cell type. Harmony-integrated clustering was repeated on each cell type independently, and clusters were removed if they were enriched for ribosomal subunit genes, mitochondrial genes, or postmortem artifact-associated immediate early genes^67^. Clusters defined solely by low UMI counts with no upregulated marker genes were also excluded. Within glial cell types, clusters present in fewer than 30% of donors were removed to exclude donor-specific artifacts. These quality-filtered data were used for all subsequent zone definition and gene expression analyses.

### KDE/Zone Overlap Analysis

Using the quality-filtered data, Leiden clustering was applied at increasing resolutions on the same sNN graph to identify between 3 and 9 clusters, without recomputing the Harmony integration or dimensionality reduction. For each individual Slide-tags dataset, we estimated a smooth two-dimensional spatial density from cell coordinates using the ks R package with a bivariate Gaussian kernel and automatic bandwidth selection, with the selected bandwidth scaled by a factor of 0.9. To define spatial territories for each zone, we computed high-density regions (HDRs) at a probability mass level of p = 0.50, representing the smallest contiguous region containing 50% of the estimated density. The HDR threshold was determined by sorting KDE values in descending order and identifying the value at which the cumulative probability mass reached p. Spatial overlap between zones was computed using binary grid masks. For each zone, a Boolean mask was constructed by thresholding the KDE at the HDR cutoff. Coverage of zone A by zone B was defined as the percentage of A’s HDR grid cells that also fell within B’s HDR. Overlaps were computed for consecutive zone pairs and summarized as the mean ± SE across donors.

### Zone centroids and coordinate alignment

For each donor, zone centroids were computed as the mean x and y coordinates of all cells per zone after removing spatial outliers. For each cell, the median distance to its k nearest neighbors (k = 10, RANN R package) was calculated. Cells whose median neighbor distance exceeded 3 standard deviations above the mean across all cells were classified as spatial outliers and excluded. A dorsal–ventral spatial axis was defined as the unit vector from the zone 1 centroid to the zone 4 centroid, and a perpendicular caudate–putamen axis was derived by 90° rotation, oriented so that zone 2 fell on the positive side.

### Other MSN zonal analysis

For striosome MSNs and D1D2 hybrid MSNs, spatial placement of unsupervised clusters was visually evaluated to identify clusters that segregated into discrete spatial zones versus those that were distributed uniformly throughout the striatum. Cells belonging to spatially segregated clusters were subset and Harmony-integrated clustering was repeated to identify 4 zones, as 5 or more zones produced substantial spatial overlap. Clustering followed the same methods described above.

### Differential expression analysis

Differential expression was performed using pseudobulk aggregation, summing raw counts per donor×zone combination using Seurat’s AggregateExpression. Genes with fewer than 50 total counts across all pseudobulk samples were removed. For each cell type, we tested each sNN-assigned zone against all remaining zones using DESeq2, including donor as a fixed-effect covariate (Love et al. 2014). Log-fold change estimates were stabilized using adaptive shrinkage (ashr) in R^68^.

### Zone-specific DEG identification

To identify genes enriched in specific zones or contiguous zone combinations, we used DESeq2 results from zones defined by unsupervised clustering (adjusted P < 0.05, log₂FC > 0.5). Genes were further required to have ≥ 20% detection within the block and ≤ 15% detection outside, with outside detection also constrained to ≤ half the inside rate. Each passing gene was assigned to its highest-FC block, with ties broken by preferring shorter blocks (fewer zones), then by adjusted P-value. For multi-zone blocks, we enforced a 20% per-zone detection floor within each constituent zone and validated block edges by requiring ≥ 2-fold enrichment of edge zone detection over pooled outside detection. Genes failing edge validation were reassigned to qualifying alternatives or removed. Finally, multi-zone block genes were classified as gradient or non-gradient using pairwise DESeq2 contrasts within the block: genes with no significant intra-block differences were classified as non-gradient, while those with identifiable high and low zone subsets were classified as gradient.

### MSN-defined compartment composition analysis

To define which MSN types were clustered and to estimate their cluster radii, we first computed the inhomogeneous pair correlation function (g_inhom(r)) for each of the 8 MSN types across 18 donors (237,717 MSN cells total, excluding one donor with low quality). The g_inhom(r) function is a non-cumulative measure that gives the relative probability of finding a neighbor at exactly distance r, controlling for macro-scale density gradients. Under the inhomogeneous Poisson null model, g_inhom(r) > 1 indicates excess neighbors (clustering) at that scale. Calculation of g_inhom(r) and all associated functions was performed using the spatstat package (v.3.5) in R.

For each donor, a convex hull of all MSN cells with a 50 µm inward buffer to reduce edge effects. Cell types with fewer than 30 cells in a given donor were excluded from analysis. Adaptive bandwidth analysis was used to estimate sigma as 500 µm for D1 Matrix, D2 Matrix, D2 StrioMat, D1D2 Hybrid, STRv NUDAP, and D1 KCNJ6; sigma = 570 µm for D1 Striosome; and sigma = 720 µm for D2 Striosome. The pair correlation function was then evaluated at r = 0 to 1000 µm in 10 µm steps using pcfinhom() and an inner exclusionary radius of 25 µm. To assess whether the observed clustering was statistically significant, we computed Monte Carlo simulation envelopes using envelope() in the spatstat package, with 199 simulations and approximately 95% confidence intervals. Simulated patterns were drawn from a background density estimated from all MSN cells at a large fixed bandwidth of 1500 µm to capture large tissue-level inhomogeneity.

Compartment boundaries were defined by density-based clustering using DBSCAN, implemented in the R package dbscan. Three compartments were independently identified: Striosome (pooling D1 and D2 Striosome together, minPts = 5), STRv NUDAP (minPts = 3), and D1 KCNJ6 (minPts = 3). For each compartment type, the neighborhood radius (eps) was calibrated per donor by sweeping eps from 20 µm to an upper bound of 2 times the radius estimated by g_inhom(r) (**Figure 1H**). At each increment of eps, convex hulls were computed around each cluster (requiring at least 3 cells) and the compositional purity was evaluated as the fraction of all MSNs falling within the hulls that belonged to the target compartment. The largest eps achieving at least 90% purity was selected for each donor; if no eps met this threshold, the smallest eps producing clusters was used. To assign the Matrix compartment from the remaining tissue regions, a fixed-radius nearest neighbor assignment was used, requiring at least 1 Matrix MSN within 150 µm and more than 2 Matrix MSNs within 250 µm of the target cell. Where compartment hulls overlapped, cells were assigned to the smallest area hull using the point.in.polygon function in the sp package.

### Spatial zone assignment of non-MSN cell types

For each donor, cells were first classified as GM or WM based on majority vote among k = 5 spatial nearest neighbors (RANN R package), using the fact that all MSNs reside in GM and that unsupervised clustering distinguishes GM and WM glial subtypes. GM cells were then assigned zone labels by majority vote among k = 15 spatial nearest neighbors drawn from D1 matrix MSN reference cells, which served as the spatial reference for zonal architecture, within a maximum distance of 750 µm.

### Cell type composition analysis

Cell type proportions were computed per donor per zone, expressed as a percentage of all cells for broad cell types or as a percentage of all neurons for MSN subtypes and interneurons. D1/D2 ratios were computed per compartment (matrix, striosome) as the ratio of D1 to D2 MSN counts per donor per zone. Zone differences were tested using paired t-tests comparing each zone (2–6) against zone 1 across donors, with Holm correction for multiple comparisons.

### Zonal DESEQ and TWI

To quantify the overall magnitude of zonal transcriptomic differences, we computed transcriptome-wide impact (TWI) for each sNN assigned zone × cell type combination by applying the univariate mode of the TRADE method (TRADEtools R package) to the shrunken log-fold change and standard error estimates from each DESeq2-computed zone-vs-all contrast ^25^. To estimate confidence intervals for transcriptome-wide impact, we used a leave-one-donor-out (LODO) jackknife approach. For each cell type, we iteratively excluded one donor at a time, re-ran the pseudobulk DESeq2 analysis for each zone-vs-all contrast on the remaining donors, and recomputed TWI from the resulting shrunken log-fold change estimates. Jackknife standard errors were calculated from the variance of the leave-one-out TWI estimates, and 95% confidence intervals were constructed around the leave-one-out mean.

### Gene set enrichment analysis and redundancy consolidation via leading-edge overlap networks

Gene set enrichment analysis (GSEA) was performed on DESeq2 differential expression results using the fgseaMultilevel algorithm ^69^. Genes were ranked by DESeq2 Wald test statistic. GO Biological Process (GO:BP) gene sets were obtained from MSigDB (v2024.1, Liberzon et al., 2015), restricted to sets with 50–230 genes in the full MSigDB annotation and further intersected with the measured gene universe. To reduce redundancy among significant gene sets (|NES| ≥ 1.0, FDR < 0.05), we constructed leading-edge (LE) overlap networks separately for positively and negatively enriched sets. For each pair of significant sets, we computed the overlap fraction as |LE_i ∩ LE_j| / min(|LE_i|, |LE_j|); an edge was drawn when this fraction exceeded 0.21. Hub gene sets (degree > 12) were iteratively removed to prevent highly connected, non-specific terms from dominating the network. On the remaining graph, connected components were identified and a single "winner" was retained per component, selected by highest degree, then lowest adjusted p-value, then highest |NES|. All pruning was performed independently for positively and negatively enriched sets to prevent cross-sign interference. Non-significant sets did not undergo consolidation.

### UCell analysis, UCell neighborhood analysis

Astrocytes were classified as high or low activity for each gene set signature (e.g., TGF-β, Smoothened) by computing per-cell UCell scores and splitting at the donor-level median. Neighborhoods were then defined for each astrocyte by identifying all cells within a 200 µm spatial radius using fixed-radius nearest neighbor search (dbscan R package). For intrinsic astrocyte expression analysis, mean expression of candidate genes was computed within high and low astrocyte groups per donor and z-scored relative to the donor-level mean and standard deviation across all astrocytes. For neighborhood expression analysis, mean expression of candidate genes was computed among neighboring cells of each cell type within high-UCell and low-UCell neighborhoods per donor, z-scored relative to the full donor-level pool of that cell type. The high−low delta for each gene was tested against zero using a one-sample t-test across donors.

### Ligand Receptor interaction analysis

Spatially resolved cell-cell interaction analysis was performed using the CellChat v2 ligand-receptor database^31^, restricted to secreted signaling and cell-cell contact interactions. For multi-subunit receptor or ligand complexes, expression was computed as the geometric mean of all subunit expression values for each cell, following the CellChat convention; cells with zero expression for any subunit received a complex expression value of zero. Six cell types were analyzed as both senders and receivers: medium spiny neurons (MSN), astrocytes, oligodendrocyte precursor cells (OPC), oligodendrocytes, microglia, and vascular cells. Genes detected in fewer than 5% of cells in all cell type by zone combinations across all six zones within a donor were excluded from analysis.

For each sender cell, an interaction score was computed as the product of the sender’s ligand expression and the trimmed mean (10% trim) of receptor expression (or receptor complex expression) among receiver cells of the specified type within a 200 um spatial radius, identified using fixed-radius nearest neighbor search (frNN; dbscan R package). Receivers in all six zones contributed to the spatial receptor potential, while sender cells were restricted to zones 1 and 4 for the zonal comparison. Scores were averaged across all sender cells within each zone per donor, and the difference (zone 1 minus zone 4) was computed as the donor-level delta for each interaction. Zone 1 versus zone 4 differences were tested using paired t-tests across 18 donors, with Benjamini-Hochberg correction for multiple testing.

To assess the robustness of identified interactions beyond statistical significance, we performed targeted permutation testing for each interaction reaching FDR < 0.05 in the paired t-test. Within each donor, zone labels (zone 1 vs. zone 4) were shuffled among sender cells of the relevant type 1,000 times, and the zone 1 minus zone 4 score difference was recomputed for each permutation to generate a donor-specific null distribution. For each donor, the observed absolute delta was compared to the 99th percentile of the absolute null distribution. To ensure directional consistency, a donor was considered robust only if (1) the observed delta exceeded the 99th percentile null threshold and (2) the sign of the donor’s delta matched the consensus direction (i.e., the sign of the cross-donor mean delta from the paired t-test). Interactions were retained in the final validated set only if this sign-concordant robustness criterion was met in at least 65% of tested donors (at least 12 of 18 donors).

### Spatial accuracy

High-confidence spatial territories were assigned using K-nearest neighbors (k=50) in physical space. A cell was labeled GM (cortical layers 1–6), WM, or non-striatal GM if >40 of its 50 neighbors shared the same zone label; otherwise it was classified as "edge" and excluded from accuracy calculations. MSN accuracy was defined by the percentage of all MSNs (excluding "STRv D1 NUDAP MSN" given their presence outside of the histologically defined striatum ^42^) located within high-confidence GM, WM, or non-striatal GM territories that were correctly placed in GM. MSNs falling in edge territories were excluded from the calculation. Astrocyte accuracy was calculated by the percentage of GM and WM astrocytes defined by unsupervised clustering (agnostic of spatial position, **Figure S4A**) that were correctly placed within GM or WM neighborhoods (>80% of k=50 neighbors with concordant GM or WM assignments).

Astrocytes falling in non-striatal GM or edge territories were excluded from the concordance calculation.

### Reference manifold construction and data integration

Reference datasets were first processed using a standardized pipeline to ensure robust feature selection and batch correction following methods used for our Slide-tags data, but implemented in Scanpy to accommodate the larger dataset size. Briefly, raw counts were normalized and log-transformed. To account for inter-donor variability, we performed batch-specific scaling, where feature scaling was applied independently within each donor. Highly variable genes (HVGs) were identified across batches, and the data was reduced to the top 2,000 HVGs. Dimensionality reduction was performed via Principal Component Analysis (PCA). To generate a shared embedding space across heterogeneous samples, we employed Harmony integration, generating a corrected low-dimensional manifold. A neighborhood graph was constructed in this integrated space, followed by Leiden clustering to define initial high-resolution cellular states.

### Hierarchical label transfer of population snRNA-seq data via MapMyCells

To utilize the complex spatial gradients and discrete molecular boundaries defining zones and combinations of zones, we implemented a hierarchical imputation strategy to impute zone labels to the population data. This approach was designed to distinguish zones by leveraging combinatorial marker expressions and gradient distinctions that might be obscured in a flat, single-level clustering analysis. Starting with the highest resolution clusters, we iteratively merged groups based on the Euclidean distance between their centroids in the Harmony embedding space. At each iteration, the two closest clusters were merged, and the resulting configuration was saved as a distinct "merge step" in the hierarchy. This produced a multi-scale taxonomic tree, ranging from broad tissue domains to highly specific sub-clusters.

Labels were transferred from the reference to the query datasets using a hierarchical implementation of the MapMyCells framework^70^. This strategy utilized the precomputed hierarchical steps to assign identities with increasing levels of specificity. At each level of the hierarchy, a richer set of differentially expressed genes could be leveraged for label transfer as opposed to a one-shot approach (forcing the method to identify differentially expressed genes without). For each node in the hierarchy, we calculated cluster-specific gene expression statistics and reference markers. To ensure the model could "break ties" between transcriptionally similar zones, we selected 30 to 100 gene markers distinguishing lineages at each branch point. Query cells were mapped onto the reference hierarchy using a bootstrapped assignment approach (1,000 iterations, bootstrap factor of 0.9). This yielded both a final zonal assignment and a confidence metric for each cell at every level of the hierarchy.

The accuracy of imputed zonal identities was validated through two independent approaches. First, label transfer was performed back onto the Slide-tags dataset using K-fold validation, and the average accuracy across the runs was computed. Second, the scaled expression of zone-defining marker genes in the Slide-tags dataset was compared with imputed population-level zones and empirically defined Slide-tags zones to confirm marker consistency. Finally, we verified that the anatomical dissection region of interest from which each population-level sample was derived corresponded to the expected imputed zonal localization.

To prepare the labeled transferred data for differential expression analysis, single-cell expression profiles were aggregated at the donor level to generate pseudobulk counts. Cell-level quality metrics were averaged per donor to create donor-specific technical covariates. The resulting profiles were filtered for lowly expressed genes and normalized using the limma-voom pipeline to model the mean-variance relationship of count data.

### Linear mixed modeling of population data with Dream

To identify zone-specific aging effects, we performed differential expression using the Dream (Differential expression for Repeated Measures) extension of the limma package, as implemented in the variancePartition framework^71^. This approach utilizes linear mixed models to increase power and control for random effects in multi-donor single-cell cohorts. Donor and pooled dissection batch were included as random intercepts to account for repeated sampling of individuals and shared backgrounds. The model included an interaction term between age and zonal identity, allowing estimation of separate aging slopes for each zone. Age was included as a continuous variable divided by 10 to report log-fold changes as a per-decade effect. The baseline age coefficient captured the aging effect in the reference zone, and zone-specific interaction coefficients captured deviations from this baseline, enabling identification of genes for which aging was accelerated, decelerated, or qualitatively different across the spatial architecture of the tissue. Absolute per-zone aging effects were computed by summing the baseline age coefficient with the corresponding zone-specific interaction term. Additional biological fixed effects included: A) zone, to regress out baseline spatial expression differences; B) genomic principal components (PC1–5) to account for population structure; and C) imputed sex. Technical fixed effects included: A) donor-averaged percentage of intronic reads; B) donor-averaged percentage of mitochondrial reads; C) fraction of ambient contamination estimated by CellBender; D) log-transformed total UMI count; E) 10x kit/chemistry version; and F) biobank source. This model was applied to cell types previously identified as zonated (D1 and D2 matrix MSNs, D1 and D2 striosome MSNs, and astrocytes). All results were corrected for multiple testing using the Benjamini-Hochberg procedure to control the false discovery rate.

### Quantifying zonal aging via transcriptome-wide impact

To quantify the aggregate transcriptomic burden of aging across zones, we applied the TRADE method (TRADEtools R package)^25^ to the dream linear mixed model results. Transcriptome-wide impact (TWI) scores were computed from the full distribution of effect sizes and standard errors from the zone-specific aging coefficients, capturing the total variance in gene expression attributable to aging within each zone without relying on arbitrary significance thresholds.

To determine whether the observed differences in aging impact between dorsal and ventral zones were statistically significant, we performed paired permutation testing. For each cell type (D1 and D2 matrix MSNs, D1 and D2 striosome MSNs, and astrocytes), we compared TWI scores between the dorsal-most and ventral-most zones. A null distribution was generated over 1,000 iterations by randomly permuting dorsal-ventral zone labels within each donor (50% swap probability), preserving donor-specific covariate structure and cell counts. For each permutation, the full differential expression pipeline (dream) and TRADE calculation were re-run to obtain permuted TWI estimates. The test statistic was defined as ΔTWI = TWI_dorsal − TWI_ventral. Significance was assessed by comparing the observed ΔTWI against the null distribution using a two-tailed test with a resolution of 0.001.

### Multivariate decomposition of zonal aging trajectories

To deconvolve heterogeneous zone-specific aging responses into interpretable biological signals, we applied consensus Independent Component Analysis (ICA) to the zone-level aging log-fold changes from the dream models. This analysis was performed on cell types with significant dorsal-ventral aging divergence (D1 matrix MSNs and astrocytes).

ICA was performed using the fastICA algorithm (parallelized, C-method) across 100 independent runs for each component count (k = 2 through 6). To identify reproducible aging patterns, we adopted a cluster-based consensus approach. For each k, mixing matrices from all 100 runs were aggregated and an absolute Pearson correlation distance matrix was computed. Components were grouped using Partitioning Around Medoids (PAM) clustering, and stability was quantified as the mean pairwise absolute correlation within each cluster. The optimal number of components was selected as the highest k at which all clusters exceeded 99% stability. Final consensus patterns were defined as the medoid of each cluster, and the consensus source matrix was derived via Moore-Penrose pseudoinverse refactorization.

Standard errors of source signal values were calculated across all runs after sign-aligning individual components to their respective cluster medoids.

To quantify the exclusivity of each gene’s association with a specific aging pattern, we computed a Signed Proportion (SP) metric, defined as the loading of gene i on pattern p divided by the sum of absolute loadings across all components. Genes were ranked by SP to identify those exclusively driving distinct zonal aging trajectories.

Gene set enrichment analysis of the independent component gene loadings was performed on SP-ranked gene lists for each component using GO Biological Process gene sets. Redundant terms were filtered using a greedy algorithm that removed terms with Jaccard similarity greater than 0.7 based on leading-edge gene overlap. To identify shared spatial aging dynamics, significant GSEA terms were compared between astrocytes and D1 matrix MSNs.

### Analyses of aging-induced zonal erosion

To analyze the dynamics of zonal markers with respect to age, we computed the average log-fold change of each zone’s markers across the age-related DE set in all zones. A two-sided Wilcoxon rank sum test was performed to identify the significance of enrichment or depletion of zonal markers in the comparisons to each zonal age-related DE set.

For each gene and each donor, between-zone variance was computed as the sample variance of the gene’s pseudobulked logCPM values across all six zones. These per-donor variances were then averaged within age-defined groups. The “young quartile” was defined as the 25th percentile of the donor age distribution, while the “old quartile” was defined as the 75th percentile of the donor age distribution.

The difference of the mean between-zone variance in the old versus the young quartiles (delta_var) was then calculated (for Figure 7I). Genes were filtered to retain 9,242 genes with mean expression in the young quartile above a logCPM value of 3. Genes with high variation in the young quartile may tend to have a negative delta_var by regression to the mean alone; to address this, we implemented a permutation test in which donors from the young and old quartiles were randomly shuffled into two groups of the same size as the original young and old groups (without replacement). The delta_var metric was then recalculated and the permutation was repeated 10,000 times to obtain an empirical p value.

To compute pairwise correlations of gene expression between zones, for each zone pair, the Pearson correlation was computed across all 9,242 genes with logCPM > 3 between pairs of each donor’s zone expression vector. The correlation between donor age (years) and the pairwise transcriptomic correlation was then computed. A positive correlation indicates that the two zones become more transcriptomically similar with age.

## Bibliography

1. Lanciego, J.L., and Obeso, J.A. (2024). Functional neuroanatomy of the normal and pathological basal ganglia. Cold Spring Harb. Perspect. Med. 10.1101/cshperspect.a041617.

2. Haber, S.N., and Knutson, B. (2010). The reward circuit: linking primate anatomy and human imaging. Neuropsychopharmacology 35, 4–26.

3. Haber, S.N. (2003). The primate basal ganglia: parallel and integrative networks. J. Chem. Neuroanat. 26, 317–330.

4. Hunnicutt, B.J., Jongbloets, B.C., Birdsong, W.T., Gertz, K.J., Zhong, H., and Mao, T. (2016). A comprehensive excitatory input map of the striatum reveals novel functional organization. Elife 5. 10.7554/eLife.19103.

5. Hintiryan, H., Foster, N.N., Bowman, I., Bay, M., Song, M.Y., Gou, L., Yamashita, S., Bienkowski, M.S., Zingg, B., Zhu, M., et al. (2016). The mouse cortico-striatal projectome. Nat. Neurosci. 19, 1100–1114.

6. Crittenden, J.R., and Graybiel, A.M. (2011). Basal Ganglia disorders associated with imbalances in the striatal striosome and matrix compartments. Front. Neuroanat. 5, 59.

7. Volkow, N.D., Koob, G.F., and McLellan, A.T. (2016). Neurobiologic advances from the brain disease model of addiction. N. Engl. J. Med. 374, 363–371.

8. Voorn, P., Vanderschuren, L.J.M.J., Groenewegen, H.J., Robbins, T.W., and Pennartz, C.M.A. (2004). Putting a spin on the dorsal-ventral divide of the striatum. Trends Neurosci. 27, 468–474.

9. Graybiel, A.M., and Matsushima, A. (2023). Striosomes and matrisomes: Scaffolds for dynamic coupling of volition and action. Annu. Rev. Neurosci. 46, 359–380.

10. Graybiel, A.M., and Ragsdale, C.W., Jr (1978). Histochemically distinct compartments in the striatum of human, monkeys, and cat demonstrated by acetylthiocholinesterase staining. Proc. Natl. Acad. Sci. U. S. A. 75, 5723–5726.

11. Saunders, A., Macosko, E.Z., Wysoker, A., Goldman, M., Krienen, F.M., de Rivera, H., Bien, E., Baum, M., Bortolin, L., Wang, S., et al. (2018). Molecular Diversity and Specializations among the Cells of the Adult Mouse Brain. Cell 174, 1015–1030.e16.

12. Stanley, G., Gokce, O., Malenka, R.C., Südhof, T.C., and Quake, S.R. (2019). Continuous and Discrete Neuron Types of the Adult Murine Striatum. Neuron. 10.1016/j.neuron.2019.11.004.

13. Gokce, O., Stanley, G.M., Treutlein, B., Neff, N.F., Gray Camp, J., Malenka, R.C., Rothwell, P.E., Fuccillo, M.V., Südhof, T.C., and Quake, S.R. (2016). Cellular Taxonomy of the Mouse Striatum as Revealed by Single-Cell RNA-Seq. Preprint, 10.1016/j.celrep.2016.06.059 https://doi.org/10.1016/j.celrep.2016.06.059.

14. Märtin, A., Calvigioni, D., Tzortzi, O., Fuzik, J., Wärnberg, E., and Meletis, K. (2019). A spatiomolecular map of the striatum. Cell Rep. 29, 4320–4333.e5.

15. van Velthoven, C.T.J., Gao, Y., Kunst, M., Lee, C., McMillen, D., Chakka, A.B., Casper, T., Clark, M., Chakrabarty, R., Daniel, S., et al. (2024). The transcriptomic and spatial organization of telencephalic GABAergic neuronal types. Neuroscience.

16. Russell, A.J.C., Weir, J.A., Nadaf, N.M., Shabet, M., Kumar, V., Kambhampati, S., Raichur, R., Marrero, G.J., Liu, S., Balderrama, K.S., et al. (2024). Slide-tags enables single-nucleus barcoding for multimodal spatial genomics. Nature 625, 101–109.

17. Hu, C., Borji, M., Marrero, G.J., Kumar, V., Weir, J.A., Kammula, S.V., Macosko, E.Z., and Chen, F. (2025). Scalable spatial transcriptomics through computational array reconstruction. Nat. Biotechnol. 10.1038/s41587-025-02612-0.

18. Johansen, N.J., Fu, Y., Schmitz, M., Dubuc, A., Kempynck, N., Wirthlin, M., Garcia, A.D., Hewitt, M., Turner, M.A., Seeman, S.C., et al. (2025). Cross-species consensus atlas of the primate basal ganglia. bioRxivorg, 2025.12.15.694496. 10.64898/2025.12.15.694496.

19. Holt, D.J., Graybiel, A.M., and Saper, C.B. (1997). Neurochemical architecture of the human striatum. J. Comp. Neurol. 384, 1–25.

20. Kubota, Y., and Kawaguchi, Y. (1993). Spatial distributions of chemically identified intrinsic neurons in relation to patch and matrix compartments of rat neostriatum. J. Comp. Neurol. 332, 499–513.

21. Glasser, M.F., and Van Essen, D.C. (2011). Mapping human cortical areas in vivo based on myelin content as revealed by T1- and T2-weighted MRI. J. Neurosci. 31, 11597–11616.

22. Gangarossa, G., Espallergues, J., Mailly, P., De Bundel, D., de Kerchove d’Exaerde, A., Hervé, D., Girault, J.-A., Valjent, E., and Krieger, P. (2013). Spatial distribution of D1R- and D2R-expressing medium-sized spiny neurons differs along the rostro-caudal axis of the mouse dorsal striatum. Front. Neural Circuits 7, 124.

23. Ren, K., Guo, B., Dai, C., Yao, H., Sun, T., Liu, X., Bai, Z., Wang, W., and Wu, S. (2017). Striatal distribution and cytoarchitecture of dopamine receptor subtype 1 and 2: Evidence from double-labeling transgenic mice. Front. Neural Circuits 11. 10.3389/fncir.2017.00057.

24. Gittis, A.H., Nelson, A.B., Thwin, M.T., Palop, J.J., and Kreitzer, A.C. (2010). Distinct roles of GABAergic interneurons in the regulation of striatal output pathways. J. Neurosci. 30, 2223–2234.

25. Nadig, A., Replogle, J.M., Pogson, A.N., Murthy, M., McCarroll, S.A., Weissman, J.S., Robinson, E.B., and O’Connor, L.J. (2025). Transcriptome-wide analysis of differential expression in perturbation atlases. Nat. Genet. 57, 1228–1237.

26. Diniz, L.P., Almeida, J.C., Tortelli, V., Vargas Lopes, C., Setti-Perdigão, P., Stipursky, J., Kahn, S.A., Romão, L.F., de Miranda, J., Alves-Leon, S.V., et al. (2012). Astrocyte-induced synaptogenesis is mediated by transforming growth factor β signaling through modulation of D-serine levels in cerebral cortex neurons. J. Biol. Chem. 287, 41432–41445.

27. Diniz, L.P., Tortelli, V., Garcia, M.N., Araújo, A.P.B., Melo, H.M., Silva, G.S.S. da, Felice, F.G.D., Alves-Leon, S.V., Souza, J.M. de, Romão, L.F., et al. (2014). Astrocyte transforming growth factor beta 1 promotes inhibitory synapse formation via CaM kinase II signaling. Glia 62, 1917–1931.

28. Rallis, A., Navarro, J.A., Rass, M., Hu, A., Birman, S., Schneuwly, S., and Thérond, P.P. (2020). Hedgehog signaling modulates glial proteostasis and lifespan. Cell Rep. 30, 2627–2643.e5.

29. Chung, K.M., Kim, H., Roque, C.G., McCurdy, E.P., Nguyen, T.T.T., Siegelin, M.D., Hwang, J.-Y., and Hengst, U. (2022). A systemic cell stress signal confers neuronal resilience toward oxidative stress in a Hedgehog-dependent manner. Cell Rep. 41, 111488.

30. Allen, N.J., Bennett, M.L., Foo, L.C., Wang, G.X., Chakraborty, C., Smith, S.J., and Barres, B.A. (2012). Astrocyte glypicans 4 and 6 promote formation of excitatory synapses via GluA1 AMPA receptors. Nature 486, 410–414.

31. Jin, S., Plikus, M.V., and Nie, Q. (2025). CellChat for systematic analysis of cell-cell communication from single-cell transcriptomics. Nat. Protoc. 20, 180–219.

32. Tan, C.X., and Eroglu, C. (2021). Cell adhesion molecules regulating astrocyte-neuron interactions. Curr. Opin. Neurobiol. 69, 170–177.

33. Murai, K.K., Nguyen, L.N., Irie, F., Yamaguchi, Y., and Pasquale, E.B. (2003). Control of hippocampal dendritic spine morphology through ephrin-A3/EphA4 signaling. Nat. Neurosci. 6, 153–160.

34. Pasterkamp, R.J. (2012). Getting neural circuits into shape with semaphorins. Nat. Rev. Neurosci. 13, 605–618.

35. Vo, T., Carulli, D., Ehlert, E.M.E., Kwok, J.C.F., Dick, G., Mecollari, V., Moloney, E.B., Neufeld, G., de Winter, F., Fawcett, J.W., et al. (2013). The chemorepulsive axon guidance protein semaphorin3A is a constituent of perineuronal nets in the adult rodent brain. Mol. Cell. Neurosci. 56, 186–200.

36. Boggio, E.M., Ehlert, E.M., Lupori, L., Moloney, E.B., De Winter, F., Vander Kooi, C.W., Baroncelli, L., Mecollari, V., Blits, B., Fawcett, J.W., et al. (2019). Inhibition of Semaphorin3A promotes ocular dominance plasticity in the adult rat visual cortex. Mol. Neurobiol. 56, 5987–5997.

37. Raz, N., Lindenberger, U., Rodrigue, K.M., Kennedy, K.M., Head, D., Williamson, A., Dahle, C., Gerstorf, D., and Acker, J.D. (2005). Regional brain changes in aging healthy adults: general trends, individual differences and modifiers. Cereb. Cortex 15, 1676–1689.

38. Fjell, A.M., Walhovd, K.B., Fennema-Notestine, C., McEvoy, L.K., Hagler, D.J., Holland, D., Brewer, J.B., and Dale, A.M. (2009). One-year brain atrophy evident in healthy aging. J. Neurosci. 29, 15223–15231.

39. Ward, R.J., Zucca, F.A., Duyn, J.H., Crichton, R.R., and Zecca, L. (2014). The role of iron in brain ageing and neurodegenerative disorders. Lancet Neurol. 13, 1045–1060.

40. Labarta-Bajo, L., and Allen, N.J. (2025). Astrocytes in aging. Neuron 113, 109–126.

41. Voorn, P., Brady, L.S., Berendse, H.W., and Richfield, E.K. (1996). Densitometrical analysis of opioid receptor ligand binding in the human striatum--I. Distribution of mu opioid receptor defines shell and core of the ventral striatum. Neuroscience 75, 777–792.

42. He, J., Kleyman, M., Chen, J., Alikaya, A., Rothenhoefer, K.M., Ozturk, B.E., Wirthlin, M., Bostan, A.C., Fish, K., Byrne, L.C., et al. (2021). Transcriptional and anatomical diversity of medium spiny neurons in the primate striatum. Curr. Biol. 31, 5473–5486.e6.

43. Bullmore, E., and Sporns, O. (2012). The economy of brain network organization. Nat. Rev. Neurosci. 13, 336–349.

44. Amemori, K.-I., Amemori, S., Gibson, D.J., and Graybiel, A.M. (2018). Striatal microstimulation induces persistent and repetitive negative decision-making predicted by striatal beta-band oscillation. Neuron 99, 829–841.e6.

45. Friedman, A., Homma, D., Gibb, L.G., Amemori, K.-I., Rubin, S.J., Hood, A.S., Riad, M.H., and Graybiel, A.M. (2015). A corticostriatal path targeting striosomes controls decision-making under conflict. Cell 161, 1320–1333.

46. Banghart, M.R., Neufeld, S.Q., Wong, N.C., and Sabatini, B.L. (2015). Enkephalin disinhibits mu opioid receptor-rich striatal patches via delta opioid receptors. Neuron 88, 1227–1239.

47. Liu, C., Goel, P., and Kaeser, P.S. (2021). Spatial and temporal scales of dopamine transmission. Nat. Rev. Neurosci. 22, 345–358.

48. Lerner, T.N., Shilyansky, C., Davidson, T.J., Evans, K.E., Beier, K.T., Zalocusky, K.A., Crow, A.K., Malenka, R.C., Luo, L., Tomer, R., et al. (2015). Intact-brain analyses reveal distinct information carried by SNc dopamine subcircuits. Cell 162, 635–647.

49. Poulin, J.-F., Caronia, G., Hofer, C., Cui, Q., Helm, B., Ramakrishnan, C., Chan, C.S., Dombeck, D.A., Deisseroth, K., and Awatramani, R. (2018). Mapping projections of molecularly defined dopamine neuron subtypes using intersectional genetic approaches. Nat. Neurosci. 21, 1260–1271.

50. Choi, E.Y., Yeo, B.T.T., and Buckner, R.L. (2012). The organization of the human striatum estimated by intrinsic functional connectivity. J. Neurophysiol. 108, 2242–2263.

51. Rieckmann, A., Johnson, K.A., Sperling, R.A., Buckner, R.L., and Hedden, T. (2018). Dedifferentiation of caudate functional connectivity and striatal dopamine transporter density predict memory change in normal aging. Proc. Natl. Acad. Sci. U. S. A. 115, 10160–10165.

52. Korkki, S.M., Johansson, J., Nordin, K., Pedersen, R., Bäckman, L., Rieckmann, A., and Salami, A. (2025). Dedifferentiation of caudate functional organization is linked to reduced D1 dopamine receptor availability and poorer memory function in aging. Imaging Neurosci. (Camb.) 3. 10.1162/imag_a_00462.

53. Dennis, N.A., and Cabeza, R. (2011). Age-related dedifferentiation of learning systems: an fMRI study of implicit and explicit learning. Neurobiol. Aging 32, 2318.e17–e30.

54. Hébert, J.M., and Fishell, G. (2008). The genetics of early telencephalon patterning: some assembly required. Nat. Rev. Neurosci. 9, 678–685.

55. Ihrie, R.A., Shah, J.K., Harwell, C.C., Levine, J.H., Guinto, C.D., Lezameta, M., Kriegstein, A.R., and Alvarez-Buylla, A. (2011). Persistent sonic hedgehog signaling in adult brain determines neural stem cell positional identity. Neuron 71, 250–262.

56. Gonzalez-Reyes, L.E., Verbitsky, M., Blesa, J., Jackson-Lewis, V., Paredes, D., Tillack, K., Phani, S., Kramer, E.R., Przedborski, S., and Kottmann, A.H. (2012). Sonic hedgehog maintains cellular and neurochemical homeostasis in the adult nigrostriatal circuit. Neuron 75, 306–319.

57. Ables, J.L., Decarolis, N.A., Johnson, M.A., Rivera, P.D., Gao, Z., Cooper, D.C., Radtke, F., Hsieh, J., and Eisch, A.J. (2010). Notch1 is required for maintenance of the reservoir of adult hippocampal stem cells. J. Neurosci. 30, 10484–10492.

58. Magnusson, J.P., Göritz, C., Tatarishvili, J., Dias, D.O., Smith, E.M.K., Lindvall, O., Kokaia, Z., and Frisén, J. (2014). A latent neurogenic program in astrocytes regulated by Notch signaling in the mouse. Science 346, 237–241.

59. Galli, S., Lopes, D.M., Ammari, R., Kopra, J., Millar, S.E., Gibb, A., and Salinas, P.C. (2014). Deficient Wnt signalling triggers striatal synaptic degeneration and impaired motor behaviour in adult mice. Nat. Commun. 5, 4992.

60. Vogel, J.W., Alexander-Bloch, A.F., Wagstyl, K., Bertolero, M.A., Markello, R.D., Pines, A., Sydnor, V.J., Diaz-Papkovich, A., Hansen, J.Y., Evans, A.C., et al. (2024). Deciphering the functional specialization of whole-brain spatiomolecular gradients in the adult brain. Proc. Natl. Acad. Sci. U. S. A. 121, e2219137121.

61. Ding, S.L., Royall, J.J., Sunkin, S.M., and Ng, L. (2016). Comprehensive cellular-resolution atlas of the adult human brain. J. Comp. Neurol. 524, 3127–3481.

62. Macosko, E.Z., Basu, A., Satija, R., Nemesh, J., Shekhar, K., Goldman, M., Tirosh, I., Bialas, A.R., Kamitaki, N., Martersteck, E.M., et al. (2015). Highly Parallel Genome-wide Expression Profiling of Individual Cells Using Nanoliter Droplets. Cell 161, 1202–1214.

63. Fleming, S.J., Chaffin, M.D., Arduini, A., Akkad, A.-D., Banks, E., Marioni, J.C., Philippakis, A.A., Ellinor, P.T., and Babadi, M. (2023). Unsupervised removal of systematic background noise from droplet-based single-cell experiments using CellBender. Nat. Methods 20, 1323–1335.

64. Kiku Ichihara,., Steven A. McCarroll Common inter-individual variation of cellular and molecular phenotypes of the human striatum. Submitted.

65. Degatano, K., Awdeh, A., Cox, R.S., 3rd, Dingman, W., Grant, G., Khajouei, F., Kiernan, E., Konwar, K., Mathews, K.L., Palis, K., et al. (2025). Warp analysis research pipelines: cloud-optimized workflows for biological data processing and reproducible analysis. Bioinformatics 41, btaf494.

66. Korsunsky, I., Millard, N., Fan, J., Slowikowski, K., Zhang, F., Wei, K., Baglaenko, Y., Brenner, M., Loh, P.-R., and Raychaudhuri, S. (2019). Fast, sensitive and accurate integration of single-cell data with Harmony. Nat. Methods 16, 1289–1296.

67. Marsh, S.E., Walker, A.J., Kamath, T., Dissing-Olesen, L., Hammond, T.R., de Soysa, T.Y., Young, A.M.H., Murphy, S., Abdulraouf, A., Nadaf, N., et al. (2022). Dissection of artifactual and confounding glial signatures by single-cell sequencing of mouse and human brain. Nat. Neurosci. 25, 306–316.

68. Stephens, M. (2017). False discovery rates: a new deal. Biostatistics 18, 275–294.

69. Korotkevich, G., Sukhov, V., Budin, N., Shpak, B., Artyomov, M.N., and Sergushichev, A. (2016). Fast gene set enrichment analysis. bioRxiv, 060012. 10.1101/060012.

70. Scott F. Daniel, Changkyu Lee, Tyler Mollenkopf, Matthew Lee, Joel Arbuckle, Elysha Fiabane, Mariano I. Gabitto, Nelson Johansen, Inkar Kapen, Andrew W. Kraft, Jane Lai, Su Ying Li, Ryan McGinty, Jeremy A Miller, Skyler Welch-Moosman, Sven Otto, Lane Sawyer, Noah Shepard, Carol L. Thompson, Andreas Tjärnberg, Jack Waters, Xingjian Zhen, Evan Macosko, Ed Lein, Lydia Ng, Hongkui Zeng, Shoaib Mufti, Zizhen Yao, Mike Hawrylycz MapMyCells: High-performance mapping of unlabeled cell-by-gene data to reference brain taxonomies. Submitted.

71. Hoffman, G.E., and Roussos, P. (2021). Dream: powerful differential expression analysis for repeated measures designs. Bioinformatics 37, 192–201.

