## Supplementary material for "Mesoscale molecular architecture of the human striatum across cell types and lifespan": Table S1

| Puck | Age | Sex | Total Cells | Median nUMI All Cells | Median nUMI MSNs |
|---|---|---|---|---|---|
| s12 | 36 | Male | 96,335 | 10,433 | 56,076 |
| s15 | 64 | Male | 98,292 | 11,919 | 55,136 |
| s16 | 66 | Male | 99,057 | 11,424 | 68,791 |
| s17 | 76 | Female | 64,844 | 12,123 | 66,366 |
| s18 | 50 | Female | 69,597 | 9,480 | 68,028 |
| s20 | 60 | Male | 46,066 | 12,410 | 20,219 |
| s21 | 56 | Male | 127,834 | 13,074 | 78,506 |
| s23 | 31 | Male | 151,068 | 19,912 | 68,513 |
| s26 | 71 | Male | 90,927 | 14,196 | 22,811 |
| s29 | 54 | Male | 98,415 | 10,967 | 42,556 |
| s31 | 45 | Male | 82,729 | 21,547 | 84,728 |
| s32 | 53 | Male | 137,759 | 7,841 | 49,782 |
| s33 | 59 | Male | 79,824 | 10,681 | 65,570 |
| s34 | 66 | Male | 83,502 | 11,946 | 48,429 |
| s35 | 69 | Male | 122,537 | 10,209 | 54,116 |
| s36 | 48 | Male | 93,376 | 9,176 | 45,324 |
| s37 | 88 | Female | 108,240 | 10,507 | 47,873 |
| s38 | 88 | Female | 94,774 | 12,896 | 38,439 |
| s5 | 44 | Male | 133,467 | 20,475 | 125,554 |
